# Structural titration reveals Ca^2+^-dependent conformational landscape of the IP_3_ receptor

**DOI:** 10.1101/2022.12.29.522241

**Authors:** Navid Paknejad, Vinay Sapuru, Richard K. Hite

## Abstract

Inositol 1,4,5-trisphosphate receptors (IP_3_Rs) are intracellular Ca^2+^-permeable cation channels whose biphasic dependence on cytoplasmic Ca^2+^ gives rise to cytosolic Ca^2+^ oscillations that regulate fertilization, cell division and cell death. Despite the critical roles of IP_3_R-mediated Ca^2+^ oscillations, the structural underpinnings of the biphasic Ca^2+^ dependence that underlies Ca^2+^ oscillations are incompletely understood. Here, we collected images of an IP_3_R with Ca^2+^ at concentrations spanning five orders of magnitude. Unbiased image analysis revealed that Ca^2+^ binding does not explicitly induce conformational changes but rather biases a complex conformational landscape consisting of resting, preactivated, activated, and inhibited states. Using particle counts as a proxy for free energy, we demonstrate that Ca^2+^ binding at a high-affinity site allows IP_3_Rs to activate by escaping a low-energy resting state through an ensemble of preactivated states. At high Ca^2+^, IP_3_Rs preferentially enter an inhibited state stabilized by a second, low-affinity Ca^2+^ binding site. Together, these studies provide a mechanistic basis for the biphasic Ca^2+^-dependence of IP_3_R channel activity.

## Introduction

Inositol-1,4,5-trisphosphate receptors (IP_3_Rs) are large, tetrameric cation channels that serve as the primary intracellular calcium (Ca^2+^) release channels in non-excitable cells. Expressed in the endoplasmic reticulum (ER), IP_3_Rs mediate the flow of Ca^2+^ from the ER into the cytoplasm and other cellular compartments where Ca^2+^ contributes to the regulation of cell division ^1^, differentiation ^2^, metabolism ^3^, migration ^4, 5^, and death ^6^. Consequently, dysregulation of IP_3_Rs is associated with numerous pathologies including cancer ^7–9^, neurological ^10, 11^, cardiac ^12^, and immune ^13^ diseases. IP_3_R activation requires nanomolar cytosolic Ca^2+^ and the second messenger IP_3_, whose production is stimulated by tyrosine kinase and G protein-coupled receptor signaling pathways ^14–20^. Notably, IP_3_Rs are inhibited by micromolar cytosolic Ca^2+^ concentrations, resulting in a biphasic Ca^2+^ dependence. The recursive nature of IP_3_R regulation by its permeant ion results in the emergent phenomenon of Ca^2+^ oscillations in cells. The Ca^2+^ dependence of both activation and inhibition are further modified by the concentration of IP_3_ as well as ATP, ER Ca^2+^ and numerous protein interaction partners ^21–23^. In this manner, IP_3_Rs integrate multiple upstream signals to tune the frequency and amplitude of Ca^2+^ oscillations to encode regulatory information for diverse cellular processes such as mitochondrial oxidative metabolism ^24^, gene expression ^25^, lymphocyte activation ^26^ and neuronal development ^27^.

Structural snapshots of IP_3_Rs in the presence and absence of regulatory ligands revealed the overall architecture of the channel and how ligands can stabilize conformational changes ^28–34^. These studies revealed that IP_3_Rs possess a transmembrane domain that resembles other 6 transmembrane (6TM) ion channels such as voltage-gated ion channels and TRP channels and a large cytosolic domain (CD) that contains all of the known regulatory ligand-binding sites and shares some homology with the Ryanodine Receptor (RyR) ^35^. When both Ca^2+^ binding sites are occupied, the pore remains closed regardless of IP_3_ binding status^29^. In contrast, a recent structure suggests that the pore opens when only one of the Ca^2+^ binding site is occupied in the presence of IP_3_ ^33^. Structures of the ligand-free (closed) and Ca^2+^- and IP_3_-bound (open) states further establish a structural basis for the activation of IP_3_Rs. However, many additional conformations have been resolved whose functional corollaries remains unclear. More broadly, the conformational landscape that enables IP_3_Rs to pivot from activation to inhibition in order to generate Ca^2+^ oscillations remains unknown. We therefore sought to establish high-resolution thermodynamic models of IP_3_R activation and inhibition using single-particle cryo-EM. By collecting images of human type 3 IP_3_R (hIP_3_R3) vitrified in a broad range of Ca^2+^ concentrations and treating particle abundance as a proxy for the relative free energy of each state, we evaluate how Ca^2+^ biases the conformational landscape of IP_3_Rs. These results establish the structural basis for IP_3_R-generated Ca^2+^ oscillations.

### Structural Ca^2+^ titration reveals conformational landscape of hIP_3_R3

To elucidate the mechanisms by which IP_3_ and Ca^2+^ together activate the channel, and high Ca^2+^ concentrations inhibit the channel, we collected transmission electron cryomicroscopic (cryo-EM) images of purified human Type 3 IP_3_ receptors (hIP_3_R3) prepared with saturating (200 µM) IP_3_, saturating (1 mM) ATP, and five Ca^2+^ concentrations spanning a physiological range from 1 nM to 10 µM (Figure 1A; Figure S1). Our cryo-EM conditions correspond to a range where hIP_3_R3 would be predicted to display a biphasic relationship between Ca^2+^ concentration and channel open probability. To track the Ca^2+^-dependence of the IP_3_R conformational landscape in an unbiased manner, we merged these datasets and performed image processing in aggregate (Figure S2; Table S1).

**Figure 1:**
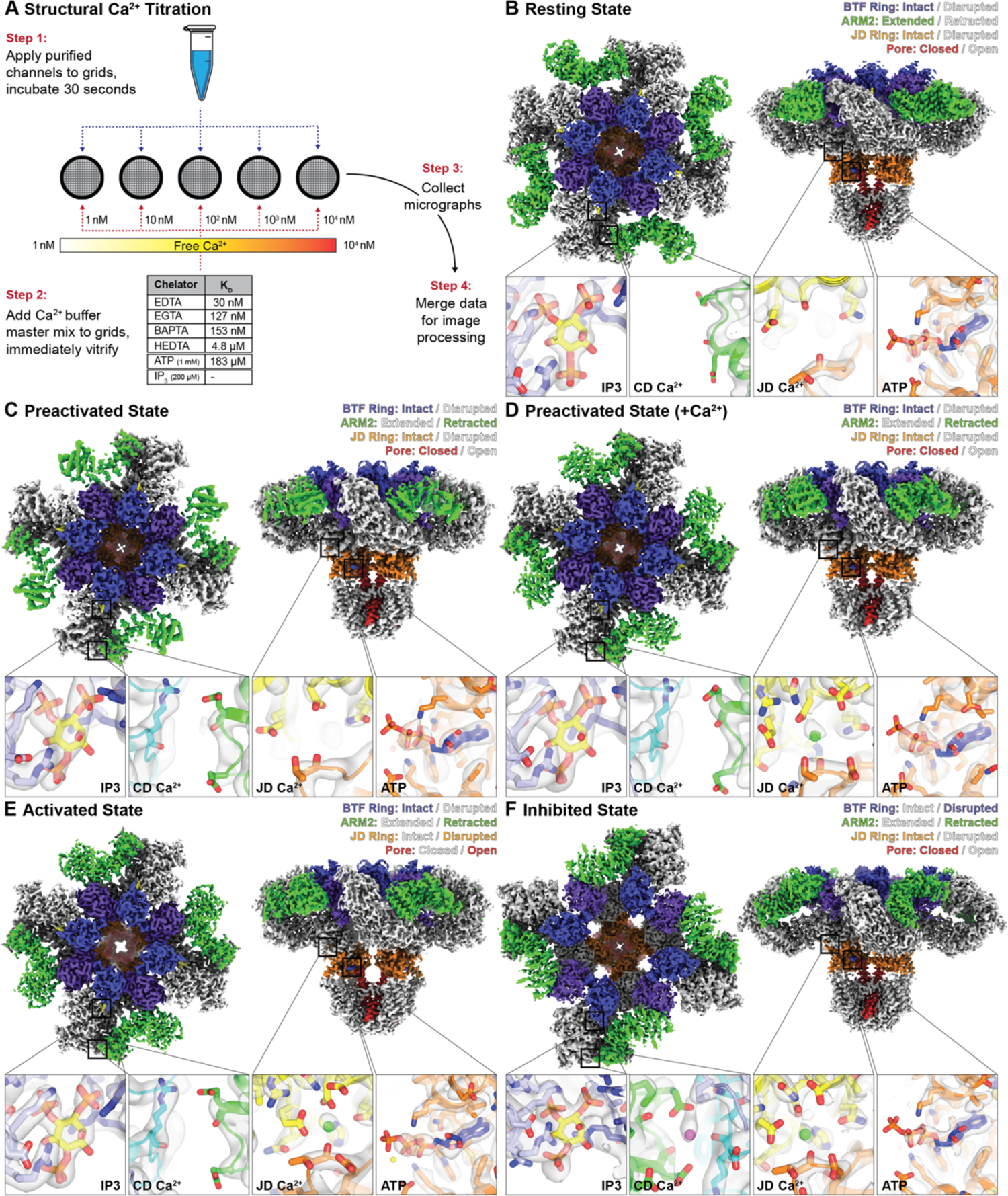
Structural Ca^2+^ titration of hIP_3_R3. **(A)** Schematic for cryo-EM Ca^2+^ titration of hIP_3_R3. **(B-F)** C4-symmetrized composite cryo-EM density maps viewed from the cytosol (left) and the side (right) with structural heuristics (top-right corner) and ligand binding status (bottom insets for IP_3_, CD Ca^2+^, JD Ca^2+^, and ATP) for the (B) resting, (C) preactivated, (D) preactivated+Ca^2+^, (E) activated, and (F) inhibited states.

Using hierarchical classification, we resolved five four-fold symmetric major states for hIP_3_R3 at resolutions up to 2.5 Å (Table S1 and S2). By relaxing the imposed C4 symmetry and computing latent representations of the conformational heterogeneity present in the remaining classes using 3D variability analysis (3DVA) ^36^, we were also able to reconstruct discrete low-abundance intermediates, including several that are asymmetric or exhibited pseudosymmetry in specific regions of the channel. Due to overlapping ligand-binding profiles of the major states and the large number of minor states, we established a heuristic describing four features of the channel that facilitate comparisons between the states as well as with existing IP_3_R structures. The features that comprise the heuristic are the beta-trefoil (BTF) ring, armadillo repeat domain 2 (ARM2), the juxtamembrane domain (JD) ring and the pore (Figure 1B-F). The most predominant of these features is the conformation of the cytosolic BTF ring, which adopts either an intact tetrameric ring structure that stabilizes the entire cytosolic domain (CD), or a disrupted state in which the CDs of the four protomers are decoupled and highly dynamic. Second is the conformation of the peripheral ARM2 domain, which can be either extended away from the rest of the CD or retracted. Third is the JD ring, which is located at the interface between the CD and the transmembrane domain (TMD) and can adopt either an intact ring structure or a disrupted, open conformation. Last is the pore, which can either be closed or open.

In the first of the major states, the BTF ring is intact, ARM2 is extended, the JD ring is intact and the pore is closed (Figure 1B; Figure S5; Movie M2; Tables S1 and S2). As this state resembles previously published ligand-free states of IP_3_Rs in various detergents (PDB: 3JAV, 6DQJ, 6MU2, 6UQK, 7LHF) and lipid environments (PDB: 7LHE) ^28–32^, we assigned this conformation as a resting state. Two similar minor states were also present that share the overall conformation of the resting state but differ slightly in the conformation of the TMD with much weaker density for the peripheral S1-S4 domain (Figure S9; Movie M7; Table S3). Due to the increased conformational heterogeneity of the TMD in these states, we assigned them as labile resting 1 and labile resting 2.

The second and third major states also have intact BTF and JD rings and a closed pore, but their ARM2 domains adopt the retracted conformation, where it is rotated towards the central linker domains (CLD) (Figures 1C-D, S7, and S8; Movies M4 and M5; Tables S1 and S2). Differentiating these two states is the presence of a non-protein density occupying the previously identified JD Ca^2+^ binding site that we assigned as a bound Ca^2+^ ion. A fourth state shares the intact BTF ring and retracted ARM2 domain with the second and third states, but its pore is open and its JD ring is disrupted (Figure 1E; Figure S6; Movie M3; Tables S1 and S2). Based on the open conformation of the pore, we assigned the fourth state as an activated state. This activated state is largely indistinguishable from a recently published structure of hIP_3_R3 with its pore in an open configuration ^33^. As the second and third states differ from the activated state only in their closed pores and intact JD rings, we assigned them as a preactivated state and a preactivated+Ca^2+^ state, respectively.

In addition to the four-fold symmetric resting and preactivated states, we also resolved classes with asymmetric CDs. In these classes, either one, two or three of the ARM2 domains adopt the retracted conformation. Together, these three classes represent a continuum of states between the resting state, where all four ARM2 domains are extended, and the preactivated state, where all four ARM2 domains are retracted, a finding we previously reported for channels in the presence of IP_3_ ^29^ (PDB: 6DQS, 6DQZ, 6DR0). While we were able to resolve structures for these states, we observed significant continuous heterogeneity among these asymmetric classes. Therefore, we combined these particles into an ensemble that we call the resting-to-preactivated transitions for quantification (Movie M6). We also observed classes with asymmetric features in the JD and TMD that otherwise resembled the resting or preactivated states. The pore in these classes has undergone movements that result in either two-fold pseudosymmetric (∼C2) or four-fold pseudosymmetric (∼C4) dilations compared to the resting state (Figure S15). We will refer to the classes with extended ARM2 domains as resting TMD transitions and those with retracted ARM2 domains as preactivated TMD transitions.

In the fifth major state, the BTF ring is disrupted, ARM2 is retracted, the JD ring is intact and the pore is closed (Figure 1E; Tables S1 and S2). A second minor population of particles sharing these features was also identified in which the channels were organized into higher-order assemblies containing two or more tetrameric channels. Notably, the interactions that mediate the assemblies are the only distinguishing feature between these two states. Otherwise, the channels adopt largely similar conformations. These two BTF ring disrupted states are reminiscent of previously published Ca^2+^-bound hIP_3_R3 structures (PDB: 6DRC, 6DR2, 6DRA, 7T3U) ^29, 33^, where BTF ring disruption confines IP_3_-induced conformational changes to the CD, so we assigned the major state as an isolated inhibited state and the minor state as an assembled inhibited state (Figure 1F; Movie M1).

### Ligand dependence of hIP_3_R3 conformations

To evaluate the relationship between ligand occupancy and conformational state, we inspected the cryo-EM maps and identified densities in the resting, preactivated, preactivated+Ca^2+^, activated and inhibited states consistent with an IP_3_ bound at the BTF2-ARM1 interface and with a Zn^2+^ and an ATP bound in the JD of all five major states (insets in Figure 1B-F). Density for IP_3_ is also present in the asymmetric subclasses that belong to the resting-to-preactivated transitions, indicating that the 200 µM IP_3_ concentration used for vitrification was sufficient to saturate the binding site ^37^, and that asymmetry of the ARM2 conformations did not arise from substoichiometric IP_3_ binding. The IP_3_-binding site is best resolved in the resting state where Arg568 on ARM1 coordinates the 1-phosphate of IP_3_ conferring a specific orientation to IP_3_ in this pocket as predicted by mutagenesis ^38^. Arg266 and Arg270 on BTF2, and Arg503, Lys507, Arg510, and Lys569 on ARM1 complete the positively charged binding site to coordinate IP_3_ (PDB: 1N4K, 3T8S, 3UJ0) ^39–41^. As observed previously ^29^, IP_3_ can bind the channel via two modes (Figure S10). Comparing the resting state to a previously published ligand-free state (PDB: 6DQJ), IP_3_ binding results in a contraction of the IP_3_-binding pocket through movement of a loop (Leu265-Ser278) on BTF2 (Figure S10A). Conversely, in the ARM2 retracted states, ARM1 tilts towards IP_3_ and the loop on BTF2 to contract the ARM1-BTF2 interface (Figure S10B-E). IP_3_ is coordinated by the same residues in both binding modes (insets in Figure 1B-F).

The Zn^2+^ ion bound in the JD is coordinated by a C_2_H_2_ zinc-finger fold formed by Cys2538, Cys2541, His2558, and His2563, where it has been observed in other IP_3_R structures ^28^ (insets in Figure 1B-F). The adenine base of the nearby ATP is buried in a hydrophobic cavity that was recently identified as an ATP-binding site that is structurally conserved with RyRs (Figure S11A-C; PDB: 7T3P, 5TAP) ^33, 42^. Specificity for adenine bases ^21, 43–45^ is imparted through the primary amine of the base forming interactions with the backbone carbonyl oxygen of His2558 and thiolate of Cys2538. The triphosphate moiety of ATP extends away from the JD with clear densities corresponding to the α and β phosphates, which are directly coordinated by Lys2152 and Lys2560, respectively. The γ-phosphate is poorly resolved and does not form direct interactions with the channel. Taken together, the coordination of ATP is consistent with both ATP and ADP having greater potentiating effects on IP_3_Rs over AMP ^21, 43, 44^.

In contrast to the saturating conditions for IP_3_ and ATP, our buffers sampled a range of Ca^2+^ concentrations that span the reported apparent affinities for both activation and inhibition of IP_3_Rs, suggesting that we might resolve a range of Ca^2+^ occupancies among the major states. To assess the Ca^2+^-dependence of each conformation, we first inspected the cryo-EM density maps near the previously identified JD and CD Ca^2+^ binding sites ^29^. In both the resting state and the preactivated state, no density peaks consistent with a bound Ca^2+^ ion were observed at either binding site (Figure 1B-F). In the preactivated+Ca^2+^ state, we observed a density peak that we assigned as a Ca^2+^ in the JD site while the CD site was unoccupied. The Ca^2+^-binding profile of the activated state is the same as the preactivated+Ca^2+^ state, with an occupied JD site and an empty CD site. Only in the inhibited state did we observe densities corresponding to Ca^2+^ in both sites. In the three JD Ca^2+^-bound states, the backbone of Thr2581 from the JD and side chains of Glu1882, Glu1946, Gln1949 from ARM3 coordinate the Ca^2+^ (Figure 3Q). The CD Ca^2+^, observed exclusively in the inhibited state, is coordinated by the backbone of Arg743 from the CLD and side chain of Glu1125 and backbone of Glu1122 from ARM2. Outside of the CD and JD sites, no densities consistent with bound Ca^2+^ ions could be identified in any of the maps. Taken together with our previous analyses of hIP_3_R3 in saturating Ca^2+^ ^29^, these data are consistent with the JD and CD sites being the primary Ca^2+^ binding sites in IP_3_Rs. Thus, in addition to their distinct global conformations, the five major states display defining ligand-binding properties. The resting and preactivated states, which bind IP_3_, ATP, and Zn^2+^, but not Ca^2+^, differ in how they coordinate IP_3_. In addition to IP_3_, ATP, and Zn^2+^, a single Ca^2+^ ion is bound to each protomer of the preactivated+Ca^2+^ and activated states, while two Ca^2+^ ions are bound to each promoter of the inhibited state.

### Ca^2+^ perturbs the energetic landscape of hIP_3_R3

Single-particle cryo-EM analysis of vitrified samples represents a near equilibrium assessment of their conformational landscape, allowing one to infer relative conformational free energy from the number of particles that populate specific structural classes ^46^. Therefore, by analyzing the effects of Ca^2+^ on the relative abundance of each hIP_3_R3 conformation or ensemble, we can assess how Ca^2+^ biases the energetic landscape of the channel to favor activation at intermediate concentrations and favor inhibition at high concentrations.

Furthermore, the Ca^2+^-dependent conformational landscape can provide additional confidence in the assignment of functional correlates to the observed conformational states (Figure 2). For example, the abundance of the putative resting state, which closely resembles the ligand-free state and shows no evidence of bound Ca^2+^ ions, is negatively correlated with the concentration of Ca^2+^. At low Ca^2+^, 45.2% of the particles adopt the resting state whereas this percentage drops to 0.7% at high Ca^2+^. Together, the two labile resting states follow a similar pattern, starting at 20.4% of the particles at 1 nM and falling to 1.2% at 10 µM. The ensemble of resting TMD transitions, comprised of the ∼C2 and ∼C4 states, is also similar, starting at 6% at 1 nM and falling to 2.4% at 10 µM.

**Figure 2:**
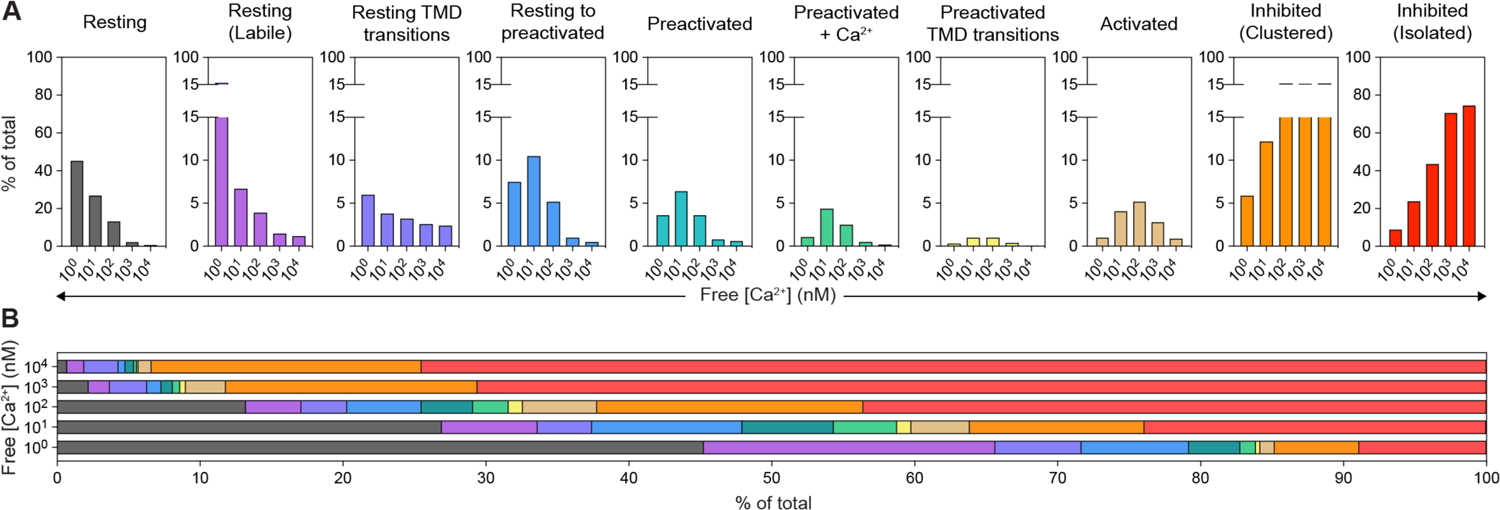
Ca^2+^-dependent conformational landscape of hIP_3_R3. **(A)** Relative percent abundance across the Ca^2+^ titration (1 nM, 10 nM, 100 nM, 1 μM, and 10 μM) for the five major states and the ensembles of minor states (denoted by *). Note that the Y-axis is truncated for several low-abundance states. **(B)** Aggregate abundances of all states across the Ca^2+^ titration.

We observed two distinct inhibited states – an isolated inhibited state and an assembled inhibited state in which several inhibited tetramers form higher-order assemblies (Figures 1E, S4 and S2). Although the states are structurally very similar with disrupted BTF rings and both Ca^2+^-binding sites being occupied, they have distinct abundance profiles with respect to Ca^2+^ concentration (Figure 2A). The abundance of the isolated inhibited channels is the inverse of the resting state i.e. positively correlated to Ca^2+^ concentration, increasing monotonically to a maximum of 74.5% at 10 µM. The assembled inhibited state channels follows the same pattern at low Ca^2+^ concentrations, increasing from 5.9% at 1 nM to a maximum of 20.1% at 100 nM. However, higher Ca^2+^ concentrations do not have any additional effect as the abundance of the assembled inhibited state plateaus between 17.6% and 20.1%. Although the structures of the tetramers in the higher-order assemblies are indistinguishable from the individual inhibited tetramers, their divergent Ca^2+^-dependence suggests that they are distinct states and that formation of higher-order assemblies may represent an alternative mechanism for achieving an inhibited state, as we will discuss later.

In contrast to the resting-like states and the inhibited states, the distribution of the preactivated-like and activated states exhibit biphasic Ca^2+^ dependencies, achieving their maximum abundance at intermediate Ca^2+^ concentrations (Figure 2A). Starting with the ensemble of resting-to-preactivated transitions, which achieve a maximum of 10.5% at 10 nM, the profiles of the preactivated, preactivated+Ca^2+^, the ensemble of ∼C2 and ∼C4 preactivated TMD transitions, and the activated state are shifted rightward to progressively higher Ca^2+^ concentrations. Apart from the activated state, the maximum abundance achieved by these states also decreases in a progressive manner, consistent with these states being progressively higher energy intermediates along a reaction coordinate extending from the resting state to the activated state. This continuum of inter-convertible states also provides a rationale for why the ensemble of resting-to-preactivated transitions and the preactivated state display a clear correlation with Ca^2+^ despite not showing evidence of binding Ca^2+^ themselves.

The abundance profile of the activated state agrees with decades of single-channel electrophysiological analyses of IP_3_Rs, showing a biphasic open probability in the presence of saturating IP_3_ and ATP with maximal activity occurring in the high nM Ca^2+^ range (Figure 2A)^19^. Moreover, the Ca^2+^-dependent conformational landscape of IP_3_Rs resolves a bipartite mechanism for this biphasic relationship with Ca^2+^ concentration. At low Ca^2+^ IP_3_Rs must escape a low-energy ARM2 extended resting state in order to activate by binding Ca^2+^ at the high-affinity JD site. At high Ca^2+^, IP_3_Rs preferentially enter a low-energy inhibited state stabilized by a second Ca^2+^ ion binding to the low-affinity CD site.

### The JD Ca^2+^ site is essential for Ca^2+^ oscillations

The multimodal regulation of IP_3_Rs, including activation and feedback inhibition by Ca^2+^, produces IP_3_R-dependent Ca^2+^ oscillations in cells ^47–50^. Structurally, we observe that Ca^2+^ binding at the JD can occur in the putative activated state, while Ca^2+^ binding at the CD site occurs only in the inhibited states. To assess the roles of these sites in cellular Ca^2+^ oscillations and to attempt to establish a functional corollary to the conformational states obtained through the structural Ca^2+^ titration, we employed a fluorescence-based Ca^2+^ imaging assay that monitors Ca^2+^ oscillations in cells. We first incubated HEK293T cells lacking all three IP_3_R isoforms (IP_3_R-null) with Cal-520-AM, a fluorogenic calcium-sensitive dye, and then stimulated intracellular IP_3_ generation by adding carbachol to the bath solution ^51^ (Figure 3B). Saturating carbachol concentrations (100 µM) were added to cells to minimize potential stimulus dependent effects on the IP_3_R response in cells ^52^. Consistent with earlier reports ^51^, no detectable changes in cytosolic Ca^2+^ were observed in IP_3_R-null cells (Figure S12P). Conversely, Ca^2+^ oscillations of two or more peaks were observed in cells transiently expressing hIP_3_R3, indicating that the construct used for structural analysis expresses a functional channel (Figure 3C-E). We assessed the temporal characteristics of the carbachol stimulated Ca^2+^ spikes in cells by aligning the initial peak of each normalized cellular trace that produced an oscillatory response (Figure 3D). For IP_3_R-null transiently expressing wild-type hIP_3_R3, the mean slope of the rising phase at the half-maximal intensity was 0.15 Fluorescence_norm_ sec^-1^. Traces were also analyzed to determine the number of peaks observed in cells showing oscillatory responses following carbachol stimulation, with cells expressing wild-type hIP_3_R3 having a median of 4 peaks/cell (Figure 3E). Finally, to calculate the time between successive Ca^2+^ spikes (inter-spike interval), we extracted traces from segmented cells, smoothed and adjusted the baseline to automatically identify peaks. For wild-type hIP_3_R3 the mean inter-spike interval was 21.7 seconds, which is within the range of times measured for endogenous IP_3_R-mediated cytosolic Ca^2+^ oscillations ^53, 54^.

Having established metrics that describe the carbachol-induced Ca^2+^ oscillations of wild-type hIP_3_R3, we next examined the effects of perturbing the Ca^2+^-binding sites. We transiently expressed hIP_3_R3 with mutations to the JD site (Glu1882Gln+Glu1946Gln), the CD site (Glu1125Gln) or both sites (Glu1125Gln+Glu1882Gln+Glu1946Gln) in IP_3_R-null cells. Robust Ca^2+^ oscillations were observed in cells expressing the CD mutant (Figure 3F-H). While the mean rising phase was similar to wild-type hIP_3_R3 (Figure 3G), the mean inter-spike interval was approximately half at 12.7 seconds (Figure S12F), suggesting that perturbing the CD site alters gating of hIP_3_R3. As the CD site is exclusively occupied in the inhibited states, our structural and functional analyses are consistent with Ca^2+^ binding at the CD site contributing to channel inhibition.

**Figure 3:**
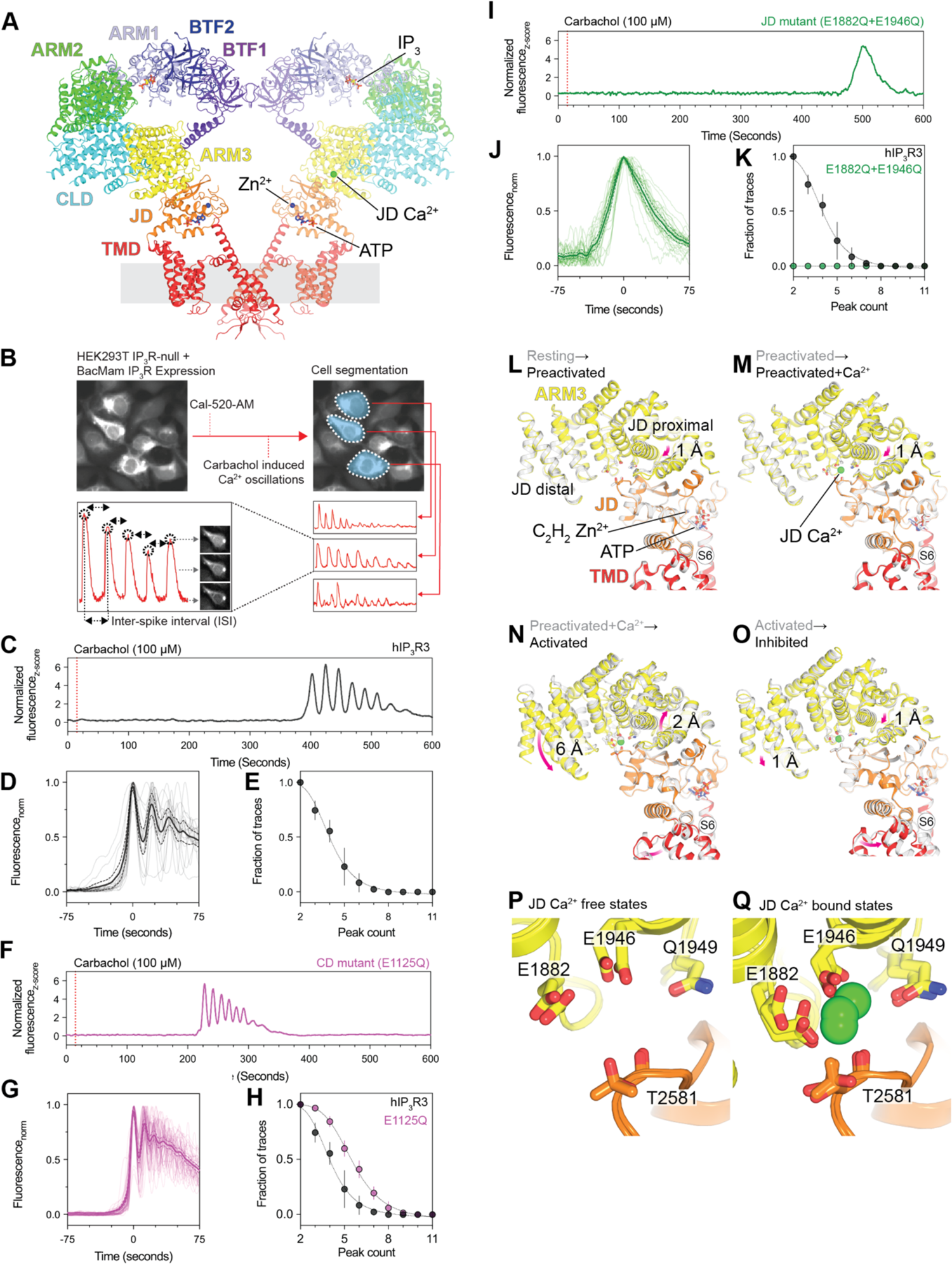
Ca^2+^ binding to the JD site has diverse effects on channel conformation. **(A)** Side view of the activated state highlighting domain architecture on the left protomer and ligand binding sites on the right protomer. Front and rear protomers removed for clarity. **(B)** Schematic describing Cal-520-AM fluorescence based Ca^2+^ imaging assay and data analysis. **(C,F,I)** Representative z-score normalized Cal-520-AM fluorescence traces recorded from cells expressing (C) hIP_3_R3, (F) CD mutant (E1125Q) and (I) JD mutant (E1882Q+E1946Q) in an IP_3_R-null background following stimulation by carbachol**. (D,G,J)** Aligned first peak of every oscillatory trace (thin lines) normalized to 1 for (D) hIP_3_R3, (G) CD mutant (E1125Q) and (J) JD mutant (E1882Q+E1946Q). Bold line represents mean and dashed lines represent 95% confidence interval. **(E,H,K)** Peak count distributions for all oscillatory traces observed for (E) hIP_3_R3, (H) CD mutant (E1125Q) and (K) JD mutant (E1882Q+E1946Q). Individual points represent mean and error bars represent S.E.M. **(L-O)** Superpositions of the ARM3-JD interface aligned by the JD for transitions from (L) resting to preactivated, (M) preactivated to preactivated+Ca^2+^, (N) preactivated+Ca^2+^ to activated, and (O) activated to inhibited. Magenta arrows highlight movements of the proximal and distal regions of the JD between states. **(P-Q)** Superpositions of the JD Ca^2+^ binding site in the (P) Ca^2+^-free states and (Q) Ca^2+^-bound states.

Unlike cells expressing wild-type channels or the CD mutant, we did not observe oscillatory responses in cells expressing either the JD mutant (Figure 3I-K) or the JD/CD double mutant (Figure S12S). Instead, we observed a single slow non-oscillatory event in both mutants that did not resemble the events seen in cells expressing the wild-type channel. The mean slope of the rising phase was 3.7 times slower for cells expressing the JD mutant (Figure 3J) and 3.0 times slower for cells expressing the JD/CD double mutant (Figure S12S) than those of cells expressing wild-type hIP_3_R3 (Figure S12T). Therefore, although perturbations to the JD site do not abolish IP_3_R-mediated Ca^2+^ release, consistent with recent electrophysiological analyses showing diminished activity of JD site mutants ^55^, the JD site is essential for ensuring the fidelity of agonist-evoked cytosolic Ca^2+^ oscillations in cells.

### Binding of the JD Ca^2+^ ion has distinct effects on channel conformation

Although Ca^2+^ binding to the JD site is required for Ca^2+^ oscillations in cells through stabilizing the activated state, it is also occupied in the preactivated+Ca^2+^ and inhibited states, both of which are closed. To gain insights into how Ca^2+^ binding stabilizes these three distinct conformations, we aligned the JD of the five major states (Figure 3L-O) to visualize the progressive changes to the JD Ca^2+^ binding site during activation and inhibition. The pair-wise comparisons reveal that large changes to the ARM3-JD interface occur exclusively during the transition from the preactivated+Ca^2+^ to the activated state: the JD-distal region of ARM3 rotates 6 Å towards the JD while the JD-proximal region shifts upwards 2 Å back to its resting state position (Figure 3N). The changes that occur during the other transitions are more subtle. For example, the transitions from resting to preactivated and from preactivated to preactivated+Ca^2+^ are each accompanied by 1 Å downward movements of the JD-proximal part of ARM3 (Figure 3L-M). Binding a second Ca^2+^ at the CD site also results in a minimal rearrangement of the ARM3-JD interface, with both the distal and proximal regions of ARM3 moving down 1 Å during the transition from the activated to inhibited state (Figure 3O). Surprisingly, despite the large conformational differences between the preactivated+Ca^2+^, activated and inhibited states, the configuration of the residues that form the JD site are nearly identical. The JD binding site appears to adopt only two conformations, a Ca^2+^-free expanded conformation in the resting and preactivated states and a Ca^2+^-bound contracted conformation in the preactivated+Ca^2+^, activated and inhibited states (Figure 3P-Q). Furthermore, we only observe stable occupancy of the JD site in the ARM2 retracted states, suggesting that the IP_3_-stabilized movement of ARM2 increases the affinity for Ca^2+^. Allosteric coupling between Ca^2+^ and IP_3_ binding is consistent with biochemical experiments suggesting that Ca^2+^ binding can increase the affinity for IP_3_ ^56, 57^, and kinetic experiments showing IP_3_ binding exposes a high-affinity Ca^2+^ binding site ^58^. In summary, although Ca^2+^ binding to the JD site results in a single, distinct conformation of the binding site, the effect of Ca^2+^ binding on channel conformation at the global level can be varied and is influenced by the global ligand-binding status of the channel.

### IP_3_ primes channel activation through a cooperative process involving ARM2 retraction

Activation of IP_3_Rs requires that all four IP_3_ binding sites be intact ^51^, suggesting that a coordinated IP_3_-mediated conformational change must occur prior to pore opening. Our previous analysis revealed that the transition between ARM2 extended and ARM2 retracted states is both IP_3_-mediated, with the retracted state only being resolved in the presence of IP_3_, and cooperative, with the four-fold symmetric extended or retracted conformations being substantially favored over the asymmetric states as opposed to a binomial distribution ^29^. We therefore hypothesized that the IP_3_ binding mode of a protomer can be sensed by its neighbors and that this communication may underlie the requirement for four intact IP_3_ binding sites. To evaluate the relationships between a single protomer and its neighbors, we performed symmetry expansion, focused refinement, and 3DVA on the CD of a single protomer, which includes the uniformly-occupied IP_3_ binding site and ARM2, for the resting-to-preactivated ensemble (Figure 4A). By calculating reconstructions for particles segmented along the primary dimension of variability, we can visualize the progression of one protomer (labeled *b* in Figure 4B-G) from the ARM2 extended conformation resolved in the resting state to the ARM2 retracted conformation of the preactivated state. In the most extended ARM2 position of the central protomer, ARM2^b^ forms two interactions with the counterclockwise protomer (labeled *a*), one with ARM1^a^, and a second with BTF1^a^ (circled *1* and *2* in Figure 4B). The transition of protomer *a* to the ARM2 retracted state is accompanied by a contraction of the ARM1-BTF2 interface around IP_3_. A consequence of this contraction is that ARM1^a^ is pulled away from ARM2^b^, disrupting one of ARM2^b^’s interprotomer interactions (Figure 4C). The diminished association with the neighboring protomer results in a more dynamic state for ARM2^b^, which manifests in weaker averaged density at its distal end (Figure 4D). The increased flexibility of ARM2^b^ destabilizes its remaining interprotomer interaction with BTF1^a^ and allows it to transiently disengage from BTF1^a^ and rotate towards CLD^b^ to adopt the retracted conformation. In the retracted conformation, ARM2^b^ establishes a new interprotomer interface with BTF1^a^ (labeled 3 in Figure 4E). ARM2^b^ retraction results in a tilt of ARM1^b^ away from ARM2 on the clockwise protomer and the entire progression repeats, enabling a cascade around the tetramer that primes the JD site for Ca^2+^ binding (Figure 4F-G).

**Figure 4:**
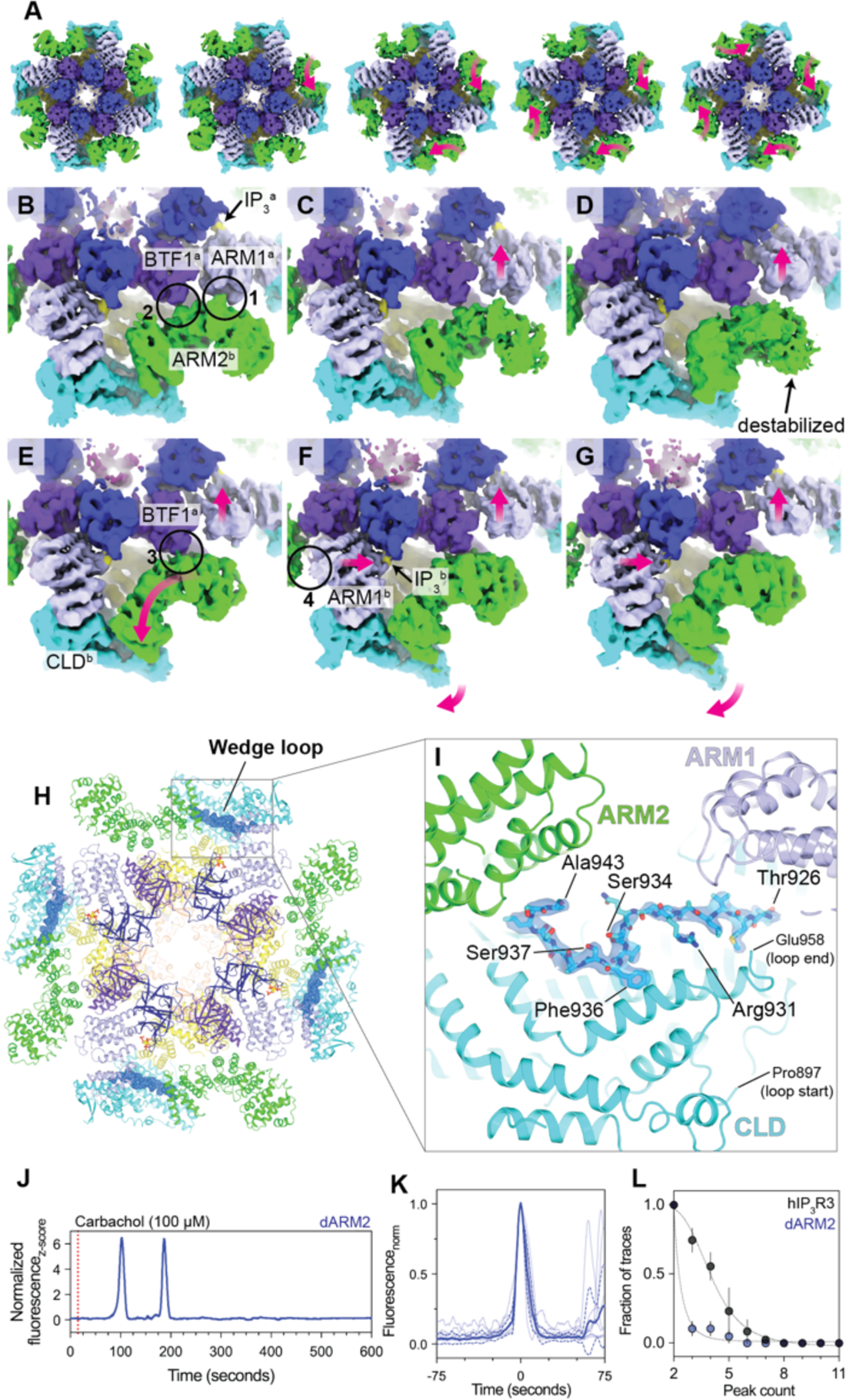
IP_3_ primes the channel for activation via a cooperative process involving ARM2. **(A)** Cryo-EM density for five states showing a distribution of ARM2 positions between the fully-extended resting state (left) and fully-retracted preactivated state (right). The three intermediates in the middle are derived from the resting-to-preactivated transitions. Magenta arrows highlight movements of the ARM2 domains compared to the preceding panel. **(B-G)** Cryo-EM density trajectory of the progression of protomer b from the extended state to the retracted state. (B) ARM2^b^ forms two interactions with the adjacent protomer at circles 1 and 2 in the ARM2 extended state. The IP_3_ bound to adjacent protomer a is highlighted. (C) The first movement is the displacement of ARM1^a^ away from ARM2^b^, which is highlighted by a pink arrow. (D) A further displacement of ARM1^a^ away is accompanied by a destabilization of the distal end of ARM2^b^. (E) ARM2^b^ is repositioned into the retracted conformation near CLD^b^ where ARM2^b^ can contact BTF1^a^ at circle 3 as ARM2^a^ continues to move towards IP3^a^. (F) Once the ARM2^b^ adopts the retracted conformation, ARM1^b^ can move towards the bound IP_3_ of protomer b, repeating the progression. This process results in torsion of the CLD^b^. (G) The movements reach their extremes in the retracted conformation. Magenta arrows highlight domain movements compared to the preceding panel. **(H)** Resting state shown as cartoon viewed from the cytosol with wedge loop shown as blue spheres. **(I)** The wedge loop occupies a cavity between ARM1, ARM2 and the CLD in the resting state. Ordered residues within the wedge loop are depicted as sticks. Cryo-EM density for the wedge loop is shown as a blue isosurface. **(J)** Representative z-score normalized Cal-520, AM fluorescence trace recorded from a cell expressing the dARM2 mutant in an IP_3_R-null background following stimulation by carbachol**. (K)** Aligned first peak of every oscillatory trace (thin lines) normalized to 1. Bold lines represent mean and dashed lines represent 95% confidence interval. **(L)** Distribution of peak counts for all oscillatory traces. Individual points represent mean and error bars represent S.E.M.

The observed continuum from a symmetric ARM2 extended resting state to a symmetric ARM2 retracted preactivated state suggests that this process is reversible despite the presence of saturating IP_3_. Consistent with the process being reversible, more particles adopt the resting state than do the ARM2 retracted preactivated and preactivated+Ca^2+^ states (Figure 2). Potentially contributing to the favorability of the ARM2 extended state is a loop between Pro897 and Glu958 of the CLD, which we call the wedge loop. In the resting state, a portion of the wedge loop, including Thr926-Ala943, inserts into a cavity surrounded by the CLD, ARM1, ARM2 and ARM3 and adopts an ordered conformation (Figure 4H-I). Compared to the resting state, ARM2 retraction in the preactivated, preactivated+Ca^2+^, activated and inhibited states is accompanied by a contraction of this cavity. Modelling the resting state conformation of Thr926-Ala943 into the ARM2 retracted states, where we observed no density for the wedge loop, reveals several steric clashes that would likely disfavor binding of the wedge loop (Figure S13D-H). To further assess the relationship between ARM2 retraction and wedge loop binding, we recalculated the ARM2 extended portion of the 3DVA trajectory of the resting-to-preactivated transitions with finer sampling. By aligning the maps based on the strength of the density for the wedge loop, we found that the flexibility of ARM2, as assessed by the local quality of the density, is inversely correlated with the strength of the wedge loop density, indicating that the presence of the wedge loop stabilizes ARM2 in the extended conformation (Figure S13J-O). Moreover, this alignment reveals how the wedge loop dissociates from its binding site in a stepwise fashion. First to dissociate are the residues surrounding Arg931, which forms a salt-bridge with Glu966 and a hydrogen bond with the hydroxyl of Tyr1067 (Figure S13C,J-L). The N- and C-terminal ends of the wedge loop become disordered in the next snapshot (Figure S13M). Phe936, which packs against Gly1073 on a helix from the CLD, is the last residue to become disordered, indicating that Phe936 is critical for the interaction (Figure S13B,N-O).

Flanking Phe936 is the conserved residue Ser934, which can be phosphorylated by protein kinase A (Figure 4I) ^59–61^. Mutation of the residue equivalent to Ser934 in hIP_3_R2 to alanine abrogates the ability of protein kinase A to sensitize hIP_3_R2 to low-level stimulation by carbachol ^62^. Modeling in a phosphorylated serine at position 934 places the phosphate group in close proximity to Ser937, potentially destabilizing the conformation of the wedge loop and weakening the critical interactions formed by Phe936, suggesting that phosphorylation of Ser934 may influence channel activity by destabilizing the resting state. Notably, the residues on and around the wedge loop described here are conserved among the three human IP_3_R isoforms, suggesting that the wedge loop may serve as regulatory motif that can influence the equilibrium between ARM2 extension and retraction and thus alters the affinity of the JD site for Ca^2+^ in all IP_3_Rs (Figures S13I and S4H).

To explore the role of the ARM2-mediated conformational changes in channel activation, we deleted the ARM2 domain (dARM2 mutant; Ala1101-Trp1586) and assessed the effects of its loss on Ca^2+^ oscillations (Figure 4J-L). Compared to cells expressing wild-type hIP_3_R3, carbachol stimulated Ca^2+^ oscillations were observed less frequently (n_WT_ = 74; n_dARM2_ = 14) in cells expressing the dARM2 mutant despite both being expressed in a similar fraction of cells (Figure 4L). Also diminished was the frequency of the Ca^2+^ spikes. The inter-spike interval was on average 4.7 times longer in cells expressing the dARM2 mutant than in cells expressing hIP_3_R3. Although the Ca^2+^ spikes were infrequent, the mean slope of the rising phase of the few responding cells was similar to that of cells expressing wild-type hIP_3_R3, suggesting that the dARM2 mutant is functional. Thus, while ARM2 is not required for activation or inhibition, its loss appears to reduce the likelihood of exceeding the threshold required for Ca^2+^ wave propagation ^49, 63^.

### Activation of hIP_3_R3 by IP_3_, Ca^2+^ and ATP

Compared to the preactivated+Ca^2+^ state, conformational changes can be observed extending from ARM3 through the JD to the TMD in the activated state (Figure 3). In both states, the JD is composed of two discontinuous segments of the polypeptide that are interwoven to connect to both the N- and C-terminal ends of the TMD (Figure 5H-I). In the preactivated+Ca^2+^ state, the four JDs assemble into a tetrameric ring structure that is also observed in the other closed states. In the activated state, the contraction of the ARM3-JD interface induces a ∼13° clockwise rigid body rotation of the JDs that disrupts the inter-JD interactions (Figure 5G) in a manner analogous to the disruption of the O-ring during RyR activation ^64^. Through its direct links to S1 and S6 (Figure 5F), the rotation of the JD alters the conformation of the central pore domain and the peripheral S1-S4 domains. The second segment of the JD, which we call JD-B comprising Cys2538-Val2611 including the Ca^2+^-binding Thr2581, is directly linked to the cytosolic end of the pore-lining S6 helix. In the activated state, rotation of the JD pulls S6 away from the center of the pore, stabilizing a 13° bend of the cytosolic end of S6 along with a ∼30° rotation about the helical axis of S6 with Gly2514 being the pivot for both. Together, the tilt and rotation of S6 reposition Phe2513 and Ile2517, which seal the ion conduction pathway in the closed states, out of the ion conduction pathway to create an open pore with a minimum radius of 4 Å (Figure 5A-E; Figure S14A-B; Movie M9-10). The repositioning of the gating residues is facilitated by rotameric switches in a manner akin to gating in Bestrophin chloride channels (Figure 5A-E) ^65^. In addition to changing the dimensions of the pore, the tilt and rotation of S6 also alter the electrostatic profile of the pore (Figure S14C-D). In the closed states, Arg2524 is oriented towards the center of the pore, creating an electropositive environment that would pose resistance to cation permeation. In the activated state, Arg2524 is rotated out of the ion conduction pathway and its place is taken by both Asp2518 and Asp2522, which render the pore electronegative and may facilitate the high cation conductance of IP_3_Rs.

**Figure 5:**
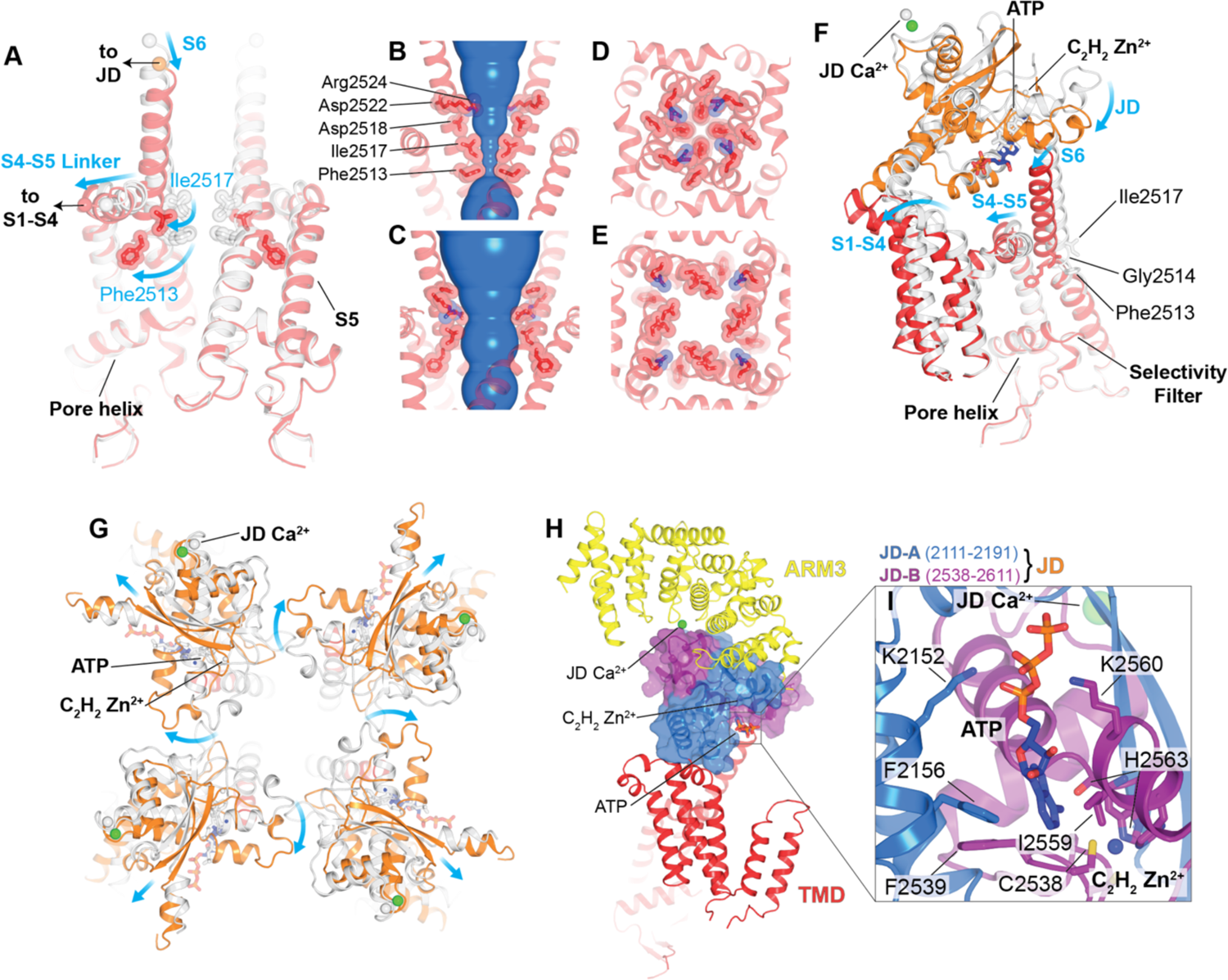
Mechanism of activation. **(A)** Superposition of the pore of the preactivated+Ca^2+^ (gray) and activated states (red), aligned by the luminal halves of S5 and S6, pore helix and selectivity filter. Front and rear protomers removed for clarity. Blue arrows highlight movement of S6, S4-S5 linker, and gating residues Phe2513 and Ile2517. Black arrows show where the pore connects to S1-S4 domain and JD. **(B-C)** HOLE diagram showing solvent-accessible surface area of conduction pathway in (B) preactivated+Ca^2+^ and (C) activated states. **(D-E)** Top view of constriction in (D) preactivated+Ca^2+^ and (E) activated states. **(F)** Comparison of TMD and JD of a single protomer of preactivated+Ca^2+^ (gray) and activated (colored) states aligned as in A. Blue arrows highlight the movements of the JD, S1-S4 bundle, S6, and the S4-S5 linker. Bending and rotation of S6 occurs at Gly2514 enabling Phe2513 and Ile2517 to repack behind the pore. **(G)** Comparison of JD ring of preactivated+Ca^2+^ (gray) and activated (colored) states viewed from the cytosol and aligned as in A. Arrows depict the movements that result in JD ring disruption during activation. **(H)** The JD (shown here in the activated state) is composed of two fragments JD-A (blue) and JD-B (purple). It is positioned between ARM3 and the TMD, and contributes to the JD Ca^2+^, ATP, and Zn^2+^ binding sites. **(I)** Inset highlights the ATP and Zn^2+^ binding sites at the interface between JD-A and JD-B.

Through its connection to S1, the first segment of the JD, which we call JD-A, can bias the conformation of the S1-S4 domain. In the closed states the peripheral S1-S4 domains adopt upright conformations that stabilize the S4-S5 linkers, which connect the S1-S4 and pore domains in domain-swapped 6TM cation channels ^66^, in a belt-like configuration that holds the pore-lining S6 helices closed (Figure 5A,F; Movie M11). In the activated state, the rotation of the JD tilts the S1-S4 domain towards the luminal side of the membrane and away from the pore (Figure 5F). This movement of the S1-S4 domains pulls the S4-S5 linkers away from the pore, thereby relaxing the belt around S6. Notably, while the S1-S4 domains remain in the upright conformation in the preactivated+Ca^2+^ state, the diminished local resolution of the S1-S4 domain suggests that Ca^2+^ binding increases the flexibility of this domain (Figure S6, S7, S8).

In addition to the fully-open activated state, our analysis identified two ensembles that contain subclasses with distorted pores that may represent snapshots of the rearrangements that occur during pore opening (Figure 6D-E). While the local resolutions of these reconstructions near the pore preclude atomic model building, comparing sections of the density maps can inform about how the pore and JDs move during gating. Among the ensemble of preactivated TMD transitions, there are two subclasses in which the conformation of the pore is altered by movements of either two pore-lining S6 helices in a ∼C2 manner or all four of the S6 helices in a ∼C4 manner. In the ∼C2 subclass, the pore-lining S6 helices from two opposing protomers shift outwards compared to their positions in the resting and preactivated states, with the remaining two helices unchanged and thus maintaining a closed pore (Figure 6D). The conformations of the S4-S5 linkers also diverge between protomers. The S4-S5 linkers of the protomers with displaced S6 helices are shifted outwards compared to the preactivated+Ca^2+^ state and are no longer in close association with the S6 helix of the neighboring protomer. This uncoupled S4-S5 linker conformation appears to be stabilized by an interaction with S1’ of the adjacent protomer (inset in Figure 6D). S1’ and S1” are two transmembrane helices unique to IP_3_Rs that are inserted between S1 and S2 of the S1-S4 domain ^29^. In all other major states, S1’ is poorly ordered and only interacts with the adjacent S1” (insets in Figure 6). In the ∼C2 transition, S1” tilts towards the pore allowing S1’ to insert underneath the S4-S5 linker of the adjacent protomer, potentially stabilizing this intermediate state. However, the precise role of S1’ and S1” in channel gating are unclear as we observed oscillatory Ca^2+^ responses in cells transduced with hIP_3_R3 in which S1’ and S1” (Glu2227-Leu2276) are deleted (Figure 6G-I). Intriguingly, while the analogous linkage between S1 and S2 is a poorly ordered acidic loop in the distantly-related RyRs ^67^, two helices preceding the S1-S4 domain occupy a similar position to S1’-S1’’ in IP_3_Rs ^42^. In the ∼C4 subclass, the S4-S5 linkers and the S6 helices of all four protomers are outwardly displaced, creating a partially dilated pore (Figure 6E). However, compared to the activated state (Figure 6F), the dilation appears to be incomplete as the cytosolic ends of S6 remain closer together. Comparing the JD in the preactivated+Ca^2+^ and activated states reveals that the JD also adopts an intermediate conformation. Whereas the JDs are both separated and rotated in the activated state, the JDs in the ∼C4 transition are only separated.

**Figure 6:**
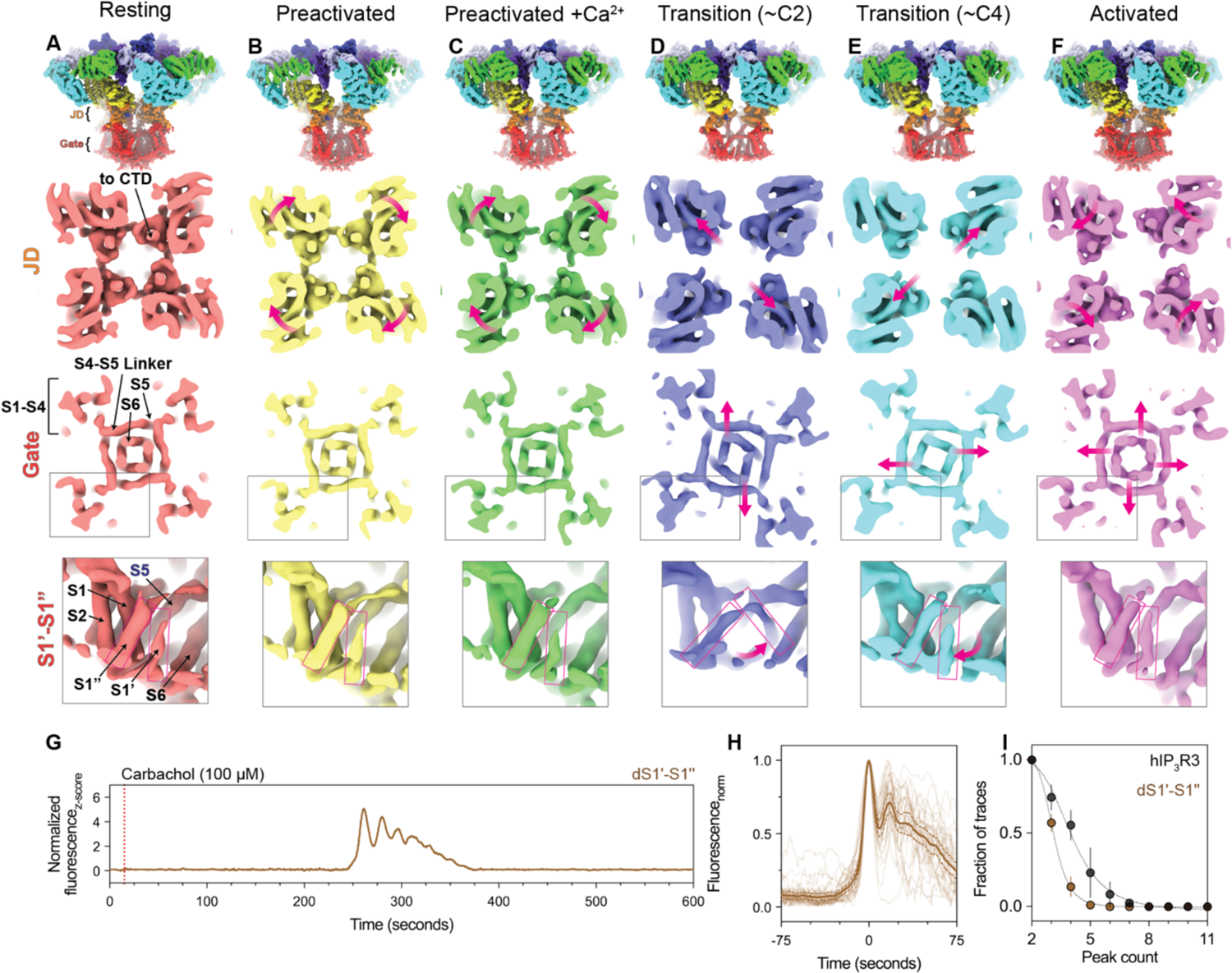
Snapshots of the conformational rearrangements in the JD and TMD that enable gating. **(A-F)** Cryo-EM density maps of the (A) resting, (B) preactivated, (C) preactivated+Ca^2+^, (D) ∼C2 preactivated TMD transition, (E) ∼C4 preactivated TMD transition, (F) activated, and (G) inhibited states, low-pass filtered to 4 Å (overall) or 7 Å (slices). **Row 1:** Overall cryo-EM density viewed from the side. **Row 2:** density slice looking from the cytosol at the height of the JD ring with magenta arrows highlight movements of the JDs. **Row 3:** density slice looking from the cytosol at the height of the gate with magenta arrows highlighting movements of the S6 helices. **Row 4:** Side view of a single S1-S4 domain with a magenta box highlighting the position of S1’-S1”. Magenta arrows denotes sequential movement of S1’ in the ∼C2 and ∼C4 preactivated TMD transition states. **(G)** Representative z-score normalized Cal-520-AM fluorescence trace recorded from a cell expressing the dS1’-S1’’ mutant in an IP_3_R-null background following stimulation by carbachol**. (H)** Aligned first peak of every oscillatory trace (thin lines) normalized to 1. Bold line represents mean and dashed lines represent 95% confidence interval. **(I)** Distribution of peak counts for all oscillatory traces. Individual points represent mean and error bars represent S.E.M.

Interpolating the ∼C2 and ∼C4 preactivated TMD transitions into a trajectory that begins with the resting state and ends with the activated state suggests a progression of JD rearrangements that facilitate gating in the pore (Figure 6A-F). First, retraction of the ARM2 domains in the preactivated state results in a clockwise rotation of the JDs which is further magnified by Ca^2+^ binding in the preactivated+Ca^2+^ state. Once Ca^2+^ is bound, the channel can sample the ∼C2 transition, where two opposing JDs shift outwards, disrupting the JD ring and leading to an outward movement of two of the four S6 helices. Then, the remaining two JDs are displaced away from the pore axis, resulting in a partial dilation of the pore. Finally, in the activated state, the JDs rotate about the helical axis of S6 to stabilize a fully-open pore where the hydrophobic gating residues Phe2513 and Ile2517 are repacked away from the permeation pathway. Notably, we do not observe any conformational changes in the pore helix or selectivity filter between the high-resolution closed and open states, indicating that the positions of Phe2513 and Ile2517 determine the gating state of the pore.

By serving as a transducer between the ligand-binding sites in the CD and the pore in the TMD, the JD is critical to regulating IP_3_R gating. At the interface between the two segments of the JD is an ATP molecule (Figure 5H-I; Figure S11). There, the adenine moiety of ATP is nestled in a hydrophobic pocket between the two segments lined by Phe2156 from JD-A and Phe2539 and Ile2559 from JD-B (Figure 5I). The phosphate groups similarly bridge the two segments of the JD with the α-phosphate coordinated by Lys2152 of JD-A and the β-phosphate coordinated by Lys2560 from JD-B. In cells, where ADP and ATP are abundant and the binding site should be predominantly occupied, ADP and ATP likely serve as molecular glue to hold the two discontinuous segments of the JD together. In the absence of ADP or ATP, Ca^2+^ binding may yield uncoupled movements of the two segments that would be a barrier to opening the pore, consistent with the prevailing model for ATP potentiation through sensitizing the channel to Ca^2+^ activation without affecting maximal open probability or high-Ca^2+^ inhibition ^19, 21^. Supporting the critical role of a rigid JD domain in channel activation, even a single cysteine-to-serine mutation at the JD Zn^2+^ binding site results in a complete loss of function without diminishing protein expression or IP_3_ affinity ^68^.

Subclasses with ∼C2 and ∼C4 distortions of the pore are also present in the ensemble of resting TMD transitions. In contrast to the preactivated TMD transitions, the JD ring remains intact in these subclasses, suggesting that conformation of the pore is not strictly coupled to that of the JD ring (Figure S15). The structural association between the JD ring and TMD in IP_3_Rs is thus weaker than the associations described between the pore and the cytosolic gating domains of other 6TM cation channels such as the BK channel (Slo1) ^69^. Alternatively, given the resemblance to the resting state, this ensemble could represent a pathway that gives rise to the previously reported ultra-low probability channel openings at very low Ca^2+^ concentrations that have been observed in the absence of IP_3_ ^70^.

### Mechanisms of high Ca^2+^ inhibition

Compared to the states with Ca^2+^ bound solely at the JD site, Ca^2+^ binding at the CD site in the inhibited state is accompanied by large conformational changes throughout the CD (Figure 1B-F). The most prominent change is the disruption of the BTF ring, which results in the CDs of the four protomers moving away from one another and towards the membrane. Despite employing the same classification approaches that resulted in identification of several other low-abundance intermediates, we did not observe any transition states between BTF ring intact and BTF ring disrupted states, suggesting that loss of a single interprotomer interaction may be sufficient to disrupt the BTF ring in a highly-cooperative fashion. Due to the presence of a second Ca^2+^ ion bound at the CD site, and because we previously demonstrated that BTF ring disruption insulates IP_3_-mediated conformational changes from the channel gate ^29^, we hypothesized that this BTF ring-disrupted conformation is the high-Ca^2+^ inhibited state of the channel. Consistent with BTF ring disruption being a key aspect of inhibition, mutations at the interface between BTF1 and BTF2 of the neighboring protomer can diminish or eliminate carbochol-induced Ca^2+^ oscillations in cells (Figure S12X). Cells expressing a Trp168Ala/Lys169Ala mutant displayed no detectable increase in cytoplasmic Ca^2+^ following carbachol stimulation, while only a single event could be observed in cells expressing a Lys169Ala mutant. These results corroborate mutagenesis experiments that predate structures of a full-length IP_3_R that yielded a graded effect on IP_3_-induced Ca^2+^ release from microsomes, with single mutations at the BTF1-BTF2 interface diminishing release compared to wild-type channels, and two or more mutations resulting in no detectable Ca^2+^ release ^71^.

Coordination of a Ca^2+^ in the CD site of the inhibited state is achieved by the N-terminal portion of the CLD and ARM2 rotating towards one another by a total of 3 Å compared to their positions in the activated state (Figure 7B,C). Through ARM1, the rotation of the CLD pulls BTF1 and BTF2 outwards, away from the BTF domains of the neighboring protomers, while the rotation of ARM2 breaks its interaction with BTF1 of the neighboring protomer. From these observations, Ca^2+^ binding to the CD site stabilizes the BTF ring disrupted conformation. However, our data cannot discern if Ca^2+^ binding at the CD site is achieved through an induced fit mechanism or through conformational selection.

**Figure 7:**
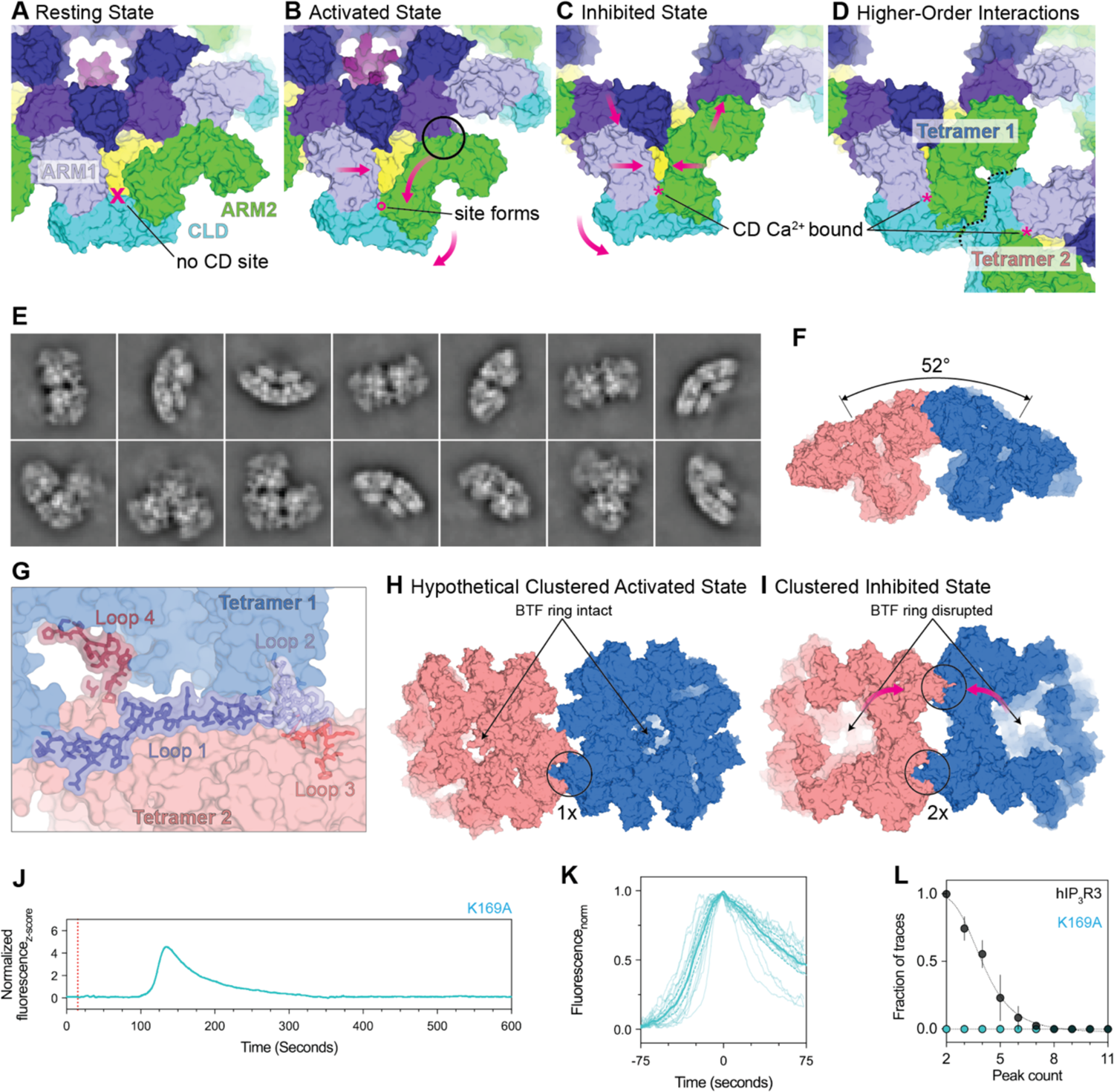
Mechanisms of high-Ca^2+^ inhibition and clustering. **(A-D)** Surface representations depicting the trajectory of a single protomer’s CD from resting to CD Ca^2+^-bound clustered inhibited states. **(A)** In the resting state, the CD Ca^2+^ binding site does not exist because ARM2 is extended away from the CLD and ARM1. Approximate location of the CD Ca^2+^ binding site in ARM2 retraced states shown as an ‘X’. **(B)** ARM2 retraction creates the CD Ca^2+^ binding site in the preactivated, preactivated+Ca^2+^, and activated states, but no Ca^2+^ is yet bound. A circle highlights an interaction between ARM2 and BTF2 of the adjacent protomer that restricts the movement of ARM2 and must be alleviated to accommodate Ca^2+^. **(C)** BTF ring disruption allows ARM2 to move further towards the CLD and bind the CD Ca^2+^ ion. **(D)** Higher-order interactions can be formed between two tetramers in a ARM2 retracted, BTF ring disrupted conformations. Dashed line represents the boundary between the two tetramers labeled Tetramer 1 and Tetramer 2 as in panel G. **(E)** Representative 2D averages of assemblies of 2-3 inhibited particles at different viewing angles. **(F)** Model of two adjacent tetramers shows a 52° angle implying a highly curved membrane environment. **(G)** Atomic model for four loops that form the higher-order interaction interface. **(H)** Modeling of a hypothetical higher-order interaction of the preactivated, preactivated+Ca^2+^, or activated state shows that steric restrictions imposed by the intact BTF ring allow only a single interaction to form between adjacent tetramers. **(I)** BTF ring disruption relieves this restriction and allows higher-order interactions to occur in a reciprocal fashion between adjacent tetramers. Magenta arrows highlight the movements of two CDs that together establish the second inter-tetramer interface. **(J)** Representative z-score normalized Cal-520-AM fluorescence trace recorded from a cell expressing the K169A mutant in an IP_3_R-null background following stimulation by carbachol**. K)** Aligned first peak of every oscillatory trace (thin lines) normalized to 1. Bold line represents mean and dashed lines represent 95% confidence interval. **(L)** Distribution of peak counts for all oscillatory traces. Individual points represent mean and error bars represent S.E.M.

While Ca^2+^ binding at the CD site stabilizes the inhibited state, Ca^2+^ oscillations, which require both activation and high-Ca^2+^ feedback inhibition ^47–50^, can be detected in cells expressing the CD site mutant (Figure 3F-H). The ability of the CD site mutant to achieve an inhibited state indicates that the CD site is not essential for inhibition and that alternative mechanisms exist. The presence of higher-order assemblies of inhibited channels suggests one potential mechanism. Although the tetramers in these assemblies are globally quite similar to the isolated inhibited channels and densities can be observed in both Ca^2+^-binding sites, they display an alternative Ca^2+^ dependence, suggesting that channels in higher-order assemblies may be functionally and structurally distinct from isolated inhibited channels.

The distinct properties of the tetramers in the higher-order assemblies may derive from the extensive state-specific interactions that stabilize the two-fold symmetric arrangement of the assembled tetramers. Four flexible linkers, which are disordered in all other states, adopt ordered conformations in assembled tetramers that contribute greatly to the two 2034 Å^2^ inter-tetramer interfaces (Figure 7G). Loop 1 (Ala1556-Asp1587), connecting the C-terminal end of ARM2 to the CLD, contributes the largest surface by snaking along the adjacent tetramer’s CLD (Figure S4A,G). Loop 2 (Pro1003-Met1023) protrudes out from the CLD to interact with Loop 3 (Phe1036-Met1044) from the adjacent tetramer (Figure S4B-C,H). Lastly, Loop 4 (Ser679-Glu690) from the CLD of the adjacent tetramer partially condenses along the inside of ARM2 (Figure S4D,I). Together, these interactions result in a 52° angle between adjacent tetramers suggesting that this architecture would be favored in highly curved membranes such as the tubular ER network (Figure 7F) ^72, 73^.

Higher-order assemblies were notably absent from investigation of the other states. By docking a model of this assembly into the other states, we discerned that the conformational restrictions imposed by an intact BTF ring allow only a single interaction to form between tetramers in the preactivated, preactivated+Ca^2+^, and activated states (Figure 7H-I). In the resting state, the extended position of ARM2 would preclude all such interactions from occurring. Together, these state-specific interactions favor the adoption of a distinct inhibited state where they can occur in a reciprocal fashion. Although there is a substantial entropic cost to these linkers adopting stable conformations, the highly ordered nature of these loops and their extensive interactions suggest that the enthalpic gains from their ordering result in an overall reduction of free energy. In the inhibited state the increased flexibility of the CD following BTF ring disruption may offset this entropic penalty. Supporting this notion, the kinetics of both elementary Ca^2+^ responses ^74^ and global Ca^2+^ oscillations ^75^ in cells exhibit strong temperature dependence.

### Flexibility of the C-terminal domain is driven by sampling acidic patches on the BTF ring

The CTD forms a four helix coiled-coil that extends through the center of the CD, connecting the JD to the BTF ring in its intact conformations (Figure S17). While functional analyses of the CTD have provided conflicting results ^68, 76, 77^, its central position led to the proposal that it may serve as an allosteric link between the IP_3_-binding sites in the CD and the pore ^30^. In hIP_3_R3, the CTD is poorly resolved due to its flexibility. Focused refinement and 3DVA revealed that a portion of the CTD of hIP_3_R3 alternatively interacts with eight negatively charged patches on the inside of the BTF ring (Figure S17A-B). While the limited resolution precludes building a model for the CTD, a conserved region of positively charged residues from Arg2654 to Arg2659 is the most likely candidate to bind to the negative patches on the BTF ring (Figure S17C). The CTD adopts two conformations which are most apparent in the activated state, interacting with four of the eight patches in either ∼C2 or ∼C4 configurations (Figure S17B; Movie M2-8), a noteworthy coincidence given the ∼C2 and ∼C4 TMD transition states. We investigated the essentiality of the CTD by truncating the channel at Leu2629 and monitoring the effects on IP_3_R-mediated Ca^2+^ oscillations. We found that while the CTD deletion (dCTD) mutant produced Ca^2+^ oscillations with a rising-phase slope that is comparable to wild-type channels, the mean inter-spike interval of 12.2 seconds is significantly shorter (Figure S12M-O). Therefore, while the CTD is not essential for channel activity, CTD deletion does alter Ca^2+^ dynamics in cells.

## Discussion

Here we defined the conformational landscape that underlies the biphasic Ca^2+^ dependence of IP_3_Rs and gives rise to IP_3_R-dependent Ca^2+^ oscillations in cells. Ordering the states based on their Ca^2+^ dependence frames a model for the ligand-dependent activation and inhibition of IP_3_Rs (Figure 8). IP_3_ generated in response to extracellular stimuli can bind to the ligand-free channel without altering its global conformation, yielding the low-energy resting state. Once bound to the resting state, IP_3_ enables the progression through the resting-to-preactivated transitions to the higher energy preactivated state, which appears to have a greater affinity for Ca^2+^. With the increased affinity, basal Ca^2+^ in the cytosol would then be able to bind to the JD site, unlocking the JD ring and favoring the transition through the ensemble of high-energy intermediate states along the trajectory to the fully-open activated state. Upon opening, IP_3_Rs release Ca^2+^ in the cytosol where it can bind to the low-affinity CD site and stabilize the inhibited state to terminate Ca^2+^ release. With IP_3_Rs closed, SERCA would be able to pump Ca^2+^ back into the ER and restore basal Ca^2+^ concentrations. As Ca^2+^ is sequestered back into the ER, Ca^2+^ can dissociate from the low-affinity CD site. When the BTF ring reforms, subsequent Ca^2+^ release events can then be initiated if IP_3_ remains abundant, resulting in regenerative Ca^2+^ oscillations.

**Figure 8:**
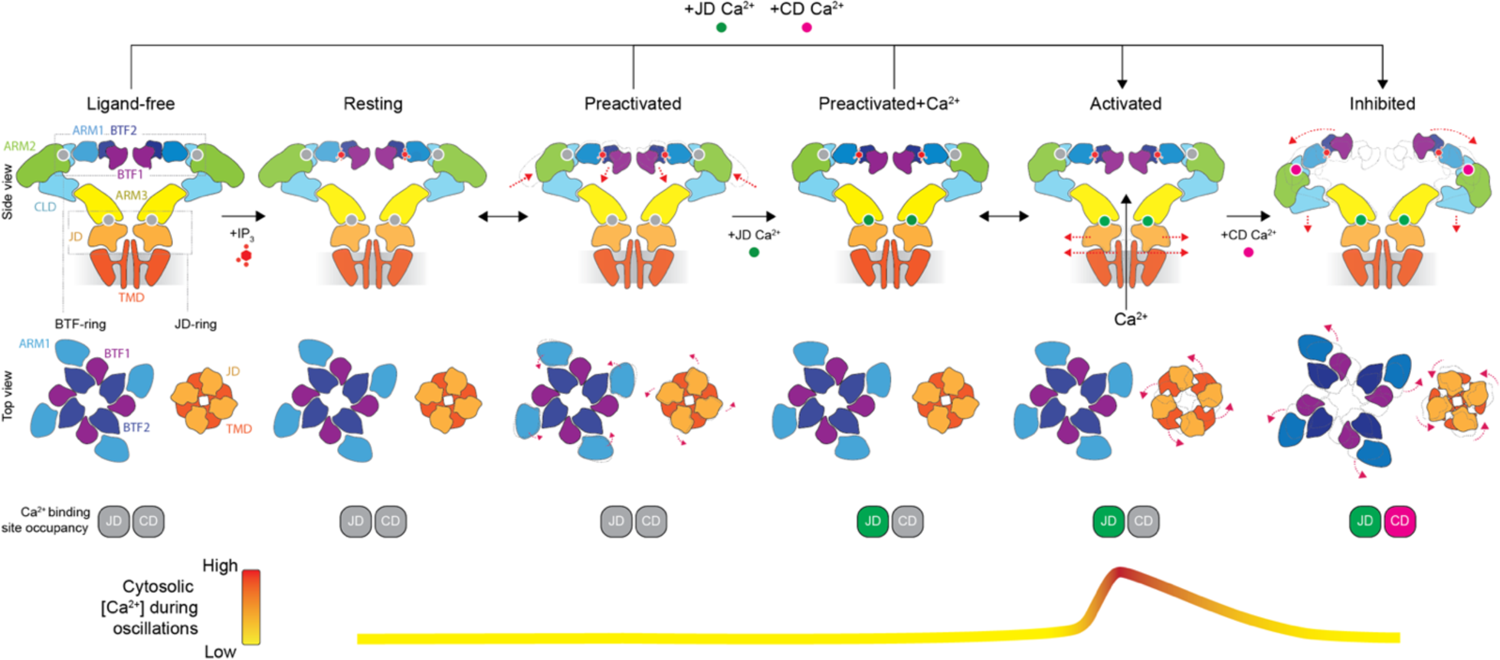
Model for biphasic regulation of IP_3_Rs by cytosolic Ca^2+^. Schematic representations depicting the mechanisms of Ca^2+^- and IP_3_-dependent activation and Ca^2+^-dependent inhibition of IP_3_Rs. **Row 1:** Side views of the major states with front and rear protomers removed for clarity. **Row 2:** Cytosolic views of BTF-ring (left) and JD-ring (right). Magenta arrows highlight movements compared to previous state. **Row 3:** Occupancy of Ca^2+^-binding sites. **Row 4:** Correspondence between conformational state and cytosolic Ca^2+^ during Ca^2+^ oscillations.

Thus, the conformational landscape of hIP_3_R3 is comprised of multiple structurally distinct closed states and seemingly only one open state. Notably, ligand binding is not sufficient to determine conformational state, as distinct states exhibit identical ligand-binding profiles. For example, the preactivated+Ca^2+^ and activated states both bind IP_3_, ATP, and Ca^2+^ at the JD site, yet the pore is closed in the preactivated+Ca^2+^ state and open in the activated state (Figure 1). Similarly, the resting and preactivated states, as well as the intermediate resting-to-preactivated states, all bind IP_3_ and ATP, but not Ca^2+^ (Figure 1). Thus, the free energy gains associated with ligand binding are insufficient to drive the ligand-induced conformational changes, such as priming and gating, to completion. Rather, ligand binding biases the conformational equilibrium to increase the favorability of the high-energy states along the trajectory to activation.

While the trajectory to activation is populated with numerous high-energy states, the resting and inhibited states serve as the lowest energy states in the low and high Ca^2+^ conditions, respectively. As the resting and inhibited states are both closed, their alternating free energy profiles contribute towards establishing the biphasic Ca^2+^ dependence of IP_3_Rs. This energetic landscape presents a highly tunable system where post-translational modifications, protein-protein interactions, membrane lipid content, and other forms of regulation can tune the balance of states to modulate activation or alter the frequency and amplitude of Ca^2+^ waves without disturbing the principal biphasic Ca^2+^ dependence of the channel. Consistent with this model, we identified several mutations that change the conformational landscape resulting in altered IP_3_R-dependent Ca^2+^ oscillation dynamics without abolishing activation or inhibition of the channel (Figures 3F-H, 4J-L, 6J-I and S12V-X). We also identified a second set of mutations that abolished Ca^2+^ oscillations, likely by removing one or more critical states from the conformational network (Figures 3I-K, 7J-L and S12R-U). Thus, our structural landscape provides a framework for understanding how diverse stimuli modulate Ca^2+^ dynamics in cells ^24, 54, 78^.

Electrophysiological analyses have demonstrated that inhibition of IP_3_R3 is highly cooperative, while activation is not ^19, 79^. These observations are consistent with the existence of multiple asymmetric states along the trajectory to activation and the complete absence of states along the trajectory to inhibition. The absence of any channels with partially disrupted BTF rings likely arises from the strain that accompanies Ca^2+^ binding. Once even a single interface in the BTF ring is disrupted, the strain throughout the channel may cause the other interfaces to be pulled apart, resulting in the inhibited state. While the CD adopts several asymmetric states in the resting-to-preactivated transitions, all of the observed states display at least two-fold pseudosymmetry in the TMD. The lack of lower symmetries in the TMD may arise from the domain-swapped arrangement of the S1-S4 domain with respect to the pore, which assures cross-protomer communication. Similar ∼C2 states have been observed for TRP channels, which share the domain-swapped 6TM fold ^80–82^. The presence of multiple transition states resolved in our analysis of hIP_3_R3 thus contrasts with two prior structural titrations of the Slo2.2 and GIRK K^+^ ion channels where intermediate states were noticeably absent and the transitions from open to closed were highly cooperative processes ^46, 83^. As electrophysiological analyses of Slo2.2 and GIRK also demonstrate this cooperativity, the correspondence between the structural and functional titrations of these three ion channels indicates that structural titrations can provide mechanistic insights into the processes that underly protein function.

It has long been appreciated that IP_3_Rs can function in higher-order assemblies, or clusters, that alter channel activity ^84^. Our studies provide structural evidence for the mechanistic underpinnings of this regulation. We observe channels in higher-order assemblies exclusively when they adopt the BTF ring disrupted inhibited state. These state-specific assemblies would allow nearby channels to transition into the inhibited state in a synchronous manner, consistent with previous analyses demonstrating that the termination of elementary IP_3_R Ca^2+^ signals produced by channels in close proximity are not completely independent ^85, 86^. Notably, formation of these higher-order assemblies requires that the ARM2 domains be in the IP_3_-stabilized retracted conformation, explaining why IP_3_ is required for cluster formation ^87, 88^. Intriguingly, the abundance of the higher-order assemblies reaches a plateau at 100 nM Ca^2+^, the same concentration at which the activated state is most abundant, suggesting that the formation of the higher-order assemblies may be primarily driven by Ca^2+^ binding to the high-affinity JD site, rather than the low-affinity CD site (Figure 2A). Ca^2+^ binding at the JD site promoting inhibition would provide an elegant failsafe mechanism to avoid excessive Ca^2+^ release and would explain how Ca^2+^ oscillations can be observed in cells expressing the CD mutant.

Altogether, our analyses show how structural titrations, the process of determining structures in the presence of varying concentrations of regulatory ligands and co-factors, can reveal how stimuli bias the conformational landscape to modulate protein function.

## Data and Code Availability

Cryo-EM maps and atomic coordinates have been deposited with the Electron Microscopy Data Bank and PDB under accession codes XXXX and EMDB-XXXX. Code is available at XXX. Summary data is available with the manuscript.

## Acknowledgements

We thank Jason de la Cruz at the Memorial Sloan Kettering Cancer Center (MSKCC) Richard Rifkind Center for cryo-EM assistance with data collection and the MSKCC High-Performance Computing (HPC) group, in particular Neeraj Harikrishnan and Jamie Cheong, for assistance with data processing. We thank Ellen Zhong for discussions about conformational heterogeneity in cryo-EM data and Elizabeth Campbell, Seth Darst, Melinda Diver and Stephen B. Long for comments on the manuscript. This work was supported by NIH NCI Cancer Center Support grant P30 CA008748 (R.K.H.), NIGMS R01-GM13230704 (R.K.H.), NCI F31-CA243235 (N.P.), the Searle Scholars Program (R.K.H.) and the Josie Robertson Investigators Program (R.K.H.).

## Author Contributions

N.P., V.S. and R.K.H. conceptualized the project and contributed to writing the manuscript. N.P. performed the bulk of cryo-EM analysis. V.S. performed the bulk of optical Ca^2+^ imaging analysis. N.P., V.S. and R.K.H. assisted each other on all experiments and analysis.

## Materials & Methods

### hIP_3_R3 expression

All constructs were N-terminally tagged with 10xHis followed by EGFP (Ca^2+^ imaging) or mVenus (cryo-EM) ^89^ followed by human rhinovirus 3C protease ^90^ cut-site and then human type 3 IP_3_R. Plasmids were transformed into DH10Bac cells to generate bacmids as described previously ^29^. 100-200 µg of purified bacmid in 1 mL water were incubated with 400 µg of 25000 MW polyethyleneimine (PEI; Polysciences Cat# 23966) in 1 mL water at 55 degC for 30-45 minutes to sterilize, then added to 50 mL of Sf9 cells at 1×10^6 cells/mL grown in suspension at 27-30 degC. The Sf9 TNMFH media was supplemented with 1% penicillin/streptomycin, 0.1% Pluronic F-68 non-ionic surfactant (Gibco Cat# 24040), and 4-8% fetal bovine serum to stabilize the virus. Virus titer was amplified to P3 and separated from cell debris by centrifugation. P3 virus was used to infect mammalian HEK293S GnTI-(ATCC CRL-3022) cells at a density of 3×10^6 cells/mL at a ratio of 50 mL virus for 800 mL cells and simultaneously stimulated with 3.75 mM valproic acid (VPA; Sigma Cat# P4543). Pellets were harvested from cells by centrifugation at 48-72 hours after infection and snap frozen.

### hIP_3_R3 purification

All surfaces, vessels, and transfer plastics were washed extensively with reverse osmosis water prior to use to minimize contaminating Ca^2+^. Membrane proteins were solubilized from 2.4 L of pelleted HEK293S GnTI-cells expressing wild-type hIP_3_R3 for 2 hours by rotation in 2% lauryl maltose neopentyl glycol (LMNG; Anatrace Cat# NG310), 150 mM sodium chloride (NaCl), 20 mM HEPES pH 7.5, 1 mM phenylmethylsulfonyl fluoride (PMSF), 2.5 µg/mL aprotinin (Sigma Cat# A1153), 2.5 µg/mL leupeptin (Alfa Aesar Cat# J61188), 10 µg/mL pepstatin A (GoldBio Cat# P-020-25), 0.5 mM 4-benzenesulfonyl fluoride hydrochloride (AEBSF; EMD Millipore Cat# 101500), and a few flakes of lyophilized deoxyribonuclease (DNAse; Worthington Biochemical Cat# LS002139). The resulting cell lysate was centrifuged at 75kxg for 40 minutes. The supernatant was incubated with sepharose-coupled GFP nanobody affinity purification beads for 4 hours with gentle agitation ^91^. The protein-GFP-bead mixture was isolated in a column, and washed with 50 mL of gel filtration buffer containing 150 mM NaCl, 50 mM Tris-HCl pH 8.0, 0.02% LMNG, and 2 mM dithiothreitol (DTT). The protein was eluted from the affinity column by cleavage with genetically modified human rhinovirus 3C protease overnight. Size exclusion chromatography was performed with a Superose 6 Increase column and the resulting protein peak was pooled and concentrated to 20 mg/mL in a 1 mL, 100 kDa MWCO concentrator (Cytiva VivaSpin Cat# 28932258).

### Structural titration sample preparation

Cryo-EM sample blotting paper contributes a significant quantity of contaminating Ca^2+^ to protein preparations. We opted to produce our own low-Ca^2+^ blotting paper by treating standard blotting paper (Ted Pella Standard VitroBot Blotting Paper Cat# 47000-100) with an extensive washing protocol. Over several days and multiple buffer exchanges, we treated with approximately 6 L of 100 µM EGTA in reverse osmosis (RO) water, then 6 L of RO water with Ca^2+^ chelating beads (BIO-RAD Chelex 100 Resin Cat#142-1253), and finally 6 L of RO water alone. The treated paper was then stacked between extensively washed glass plates and subjected to vacuum for 24 hours to remove moisture and resume a flat shape. The treated filter paper is predicted to contain substantially less than 1 mM contaminating Ca^2+^ (predicted starting condition of blotting paper ^29^) and 100 µM residual EGTA (first wash condition).

To further control our sample Ca^2+^ concentrations, we engineered a Ca^2+^ chelating cocktail. By combining 2 mM each of EDTA (Kd 30 nM), EGTA (Kd 127 nM), BAPTA (Kd 153 nM), HEDTA (Kd 4.8 µM) with 1 mM of ATP (Kd 183 µM), we calculate that our buffer ensures a semi-log-linear relationship between free and total Ca^2+^ from 1 nM to 300 µM ^92^. The least well-controlled range for free Ca^2+^ was between 1 nM and 10 nM requiring addition of 864 µM total Ca^2+^, and the largest was between 10 µM and 100 µM, requiring addition of 2.0 mM total Ca^2+^. Thus, our total contaminating Ca^2+^ must be greater than 864 µM to generate a maximum 1-log-fold error in our target free Ca^2+^ across the entire titratable range, ensuring that we maintain the semi-log-linear relationship between free and total Ca^2+^ despite contaminating Ca^2+^. To minimize the impact of widely varying kinetic properties of the chelators, we generated a pre-mixed 5X solution containing 10 mM of each chelator, 5 mM ATP, 1 mM IP_3_, and 2.5 mM fluorinated fos-choline-8 (Anatrace Cat# F300F), a detergent that does not interact with hydrocarbons, to protect the protein from the air-water interface. Sensitivity analysis using MaxChelator (https://somapp.ucdmc.ucdavis.edu/pharmacology/bers/maxchelator/webmaxc/webmaxcE.htm) revealed that inaccurate pH was the largest contributor to deviations from the predicted free Ca^2+^, and thus we carefully adjusted all solutions to pH 8, and added an additional 50 mM Tris pH 8.0 to the master mix. CaCl_2_ and MgCl_2_ were added in varying quantities to generate the desired free Ca^2+^ concentration and a constant 3 mM free Mg^2+^ concentration. During grid preparation, 3.2 µL of purified protein was added to the grid and incubated for 30 seconds, after which we added 0.8 µL of the ligand master mix directly to the droplet on the grid, immediately blotted with our low-Ca^2+^ blotting paper for 2 seconds, then plunge-frozen using a ThermoFisher Vitrobot Mark IV. Since the Ca^2+^ and chelators are premixed, the free Ca^2+^ is at equilibrium in the master mix, and pipetting error when adding to the protein on the grid will have no effect on free Ca^2+^. The only deviations due to pipetting error would be [IP_3_] and [ATP], both of which are above saturating concentrations and so we assume those to be inconsequential for this analysis. The final grid conditions have varying free Ca^2+^, but constant 200 µM IP_3_, 1 mM ATP, 3 mM free Mg^2+^, 1.6 mM dithiothreitol (DTT), 2 mM EDTA, 2 mM EGTA, 2 mM BAPTA, 2 mM HEDTA, 50 mM Tris pH 8.0, 120 mM NaCl, 500 µM fluorinated fos-choline-8, and 159 µM LMNG.

### Cryo-EM data collection, analysis and model building

Images were collected at 0.826 Å/px magnification on an FEI Krios with Gatan K3 detector at 15 e^-^/pix/sec with 3 sec exposure (0.05 sec/frame) for a total dose of 66 e^-^/Å^2^ in automated fashion using SerialEM ^93, 94^. Five datasets were collected during the same session for each Ca^2+^ concentration on a series of grids that were prepared sequentially resulting in 637 movies at 1 nM, 2150 movies at 10 nM, 6126 movies at 100 nM, 1372 movies at 1 µM, and 3136 movies at 10 µM. A sixth dataset of 4312 movies collected at nominal 100 nM free Ca^2+^ from a grid prepared later in the sequence was collected as a technical replicate to assess experimental error (Figure S16A).

All movies were combined and processed starting in CryoSparc Live v3.3.1 for motion correction, CTF estimation, and bias-free autopicking at a rate of 380 picks/micrograph with a gaussian blob of dimensions between 166 and 240 Å, corresponding to the smallest and largest diameter of the known conformational states of IP_3_Rs. Thus, all of the following classification decisions were made in aggregate and without any *a priori* knowledge of the dataset from which particle subsets were derived. The over-picked particle stack was extracted in a 512 box and subjected to iterative CryoSparc v3.3.1 Heterogeneous Refinement ^95^ with four references corresponding to the resting, activated, inhibited, and a single consensus average of the preactivated +/- Ca^2+^ states. These references were previously determined from the combined data using traditional single-particle approaches. The remaining eight classes were pure noise decoy references generated by randomly sampling a very small number of particles via CryoSparc v3.3.1 Ab-Initio without alignment. The decoy references attract false positives, while the four high-resolution references attract true positives. These references were used for all classifications described herein.

After several rounds of “decoy” classification, the particle stack went from 7.8M particles to 1.7M particles, with 351k, 117k, 145k, and 1045k residing in the classes obtained from the resting, preactivated, activated, and inhibited references respectively. 2D classification of the discarded classes confirmed that no unintentional removal of true positives occurred. At this stage, each stack was independently subjected to an additional iteration of classification to allow fine separation of states whereby the non-self references attract particles away from the self-identifying class in cases where the particles deviate from the consensus state in subtle ways. This resulted in six classes that are depicted in the second tier of the cryo-EM workflow figure (Figure S2), with classes that refined to worse than 7 Å being discarded as junk or damaged particles.

Each of these six stacks were refined enforcing C4 symmetry to improve signal for reference-based corrections prior to Bayesian Polishing in Relion v3.1.3 ^96^. At this stage, optical groups were separated and both local and global CTF parameters were optimized in CryoSparc v3.1.1 during Non-Uniform Refinement ^97^ procedures. Due to the very large number of optical groups, it was found that the fourth-order terms of spherical aberration and tetrafoil ^98, 99^ were not being fit accurately in some groups, and hence we did not fit these terms. In aggregate the per particle, per micrograph, and per optical group corrections resulted in improvements for the resting-like stack with strong TMD density (231k particles; 3.5 Å to 2.7 Å), resting-like stack with weak TMD density (108k particles; 3.9 Å to 3.3 Å), preactivated-like stack with weak CD density (83k particles; 4.0 Å to 3.6 Å), activated-like stack (65k particles; 3.7 Å to 3.1 Å), preactivated-like stack with weak TMD density (76k particles; 3.8 Å to 3.2 Å), and inhibited-like stack (1045k particles; 3.2 Å to 2.5 Å). These stacks were subjected to one final round of classification revealing the five primary C4 symmetric states called resting (192k particles; 2.8 Å), preactivated (47k particles; 3.7 Å), preactivated+Ca^2+^ (31k particles; 3.6 Å), activated (56k particles; 3.1 Å), and inhibited (917k particles; 2.5 Å) states. Additionally, we identified several heterogeneous conformational ensembles that will be discussed after the five C4 symmetric states.

We further improved the C4 symmetric states by performing C4 symmetry expansion and local refinement to correct for subtle local asymmetries in the particles. We used a model to precisely delineate masks surrounding modular units that flex and move in unison: (1) a mask containing a single chain from the tetramer (2) the entire cytosolic domain consisting of residues 1-1697 from a single chain (3) BTF1, BTF2, and ARM1 consisting of residues 1-664 from a single chain (4) CLD, ARM3 consisting of residues 665-1100 and 1586-2074 from a single chain (5) ARM2 consisting of residues 1101-1586 from a single chain (6) TMD, JD consisting of residues 2111-2611 from a single chain. The masks generated from these models were dilated by 4 pixels and a cosine soft-edge was applied for 40 pixels, thereby avoiding ringing and mask artifacts that occur when converting hard edges in real-space to reciprocal space. Therefore, this mask retains 100% of the information at ∼3 Å away from the model, and 50% of the information at ∼25 Å away from the model. CryoSparc v3.3.1 Local Refinement resulted in resolutions ranging from 2.5 Å (TMD/JD) to 3.3 Å (ARM2) for the resting state, 3.6 Å (TMD/JD) to 6.5 Å (ARM2) for the preactivated state, 3.3 Å (BTF1/BTF2/ARM1) to 4.2 Å (ARM2) for the preactivated+Ca^2+^ state, 2.9 Å (BTF1/BTF2/ARM1) to 3.3 Å (ARM2) for the activated state, and 2.5 Å (TMD/JD) to 3.4 Å (BTF1/BTF2/RM1) for the inhibited state (Figure S3, S5, S6, S7, S8). In some highly-heterogeneous cases, the local refinements were subjected to a procedure that will be described in the treatment of the conformational ensembles to improve the resolution (e.g. BTF1/BTF2/ARM1 in the inhibited state).

The local refinements were independently subjected to Phenix v1.20.1-4487 Resolve Cryo-EM ^100^ guided only by experimental density (no model) and employing a lenient mask that contains all proteinaceous and detergent micelle density, an approach we have used previously ^101–103^. As part of the procedure, the final maps are sharpened using a half-map derived factor. The resulting density modified and sharpened maps were cropped to a single chain and used for iterative model building using coot ^104^, ISOLDE ^105^ and composite map generation using a 20-residue sliding window cross-correlation (Phenix v1.20.1-4487 Combine Focused Maps) ^106^, which we found to produce artifact-free maps when compared to Chimera’s ‘vop maximum’ command ^107^. For the highest resolution composites (resting, activated, and inhibited) the density-modified local refinements were super-sampled prior to composite generation to aid interpretation of ligands, ions, and waters. Inspection of the resulting composite maps showed that they were free of model-based overfitting, for example density for ions, ligands, and lipids remain intact despite being removed from the input model. The final models were refined against the composite map with Phenix v1.20.1-4487 Real-Space Refinement ^108^.

The remaining classes represent highly-heterogeneous conformational ensembles that we interrogated via 3D variability analysis (3DVA) ^36^. We relaxed our assumptions about symmetry by performing C4 symmetry expansion on each class. For the resting-like ensemble with weak CD density, resting-like ensemble with weak TMD density, preactivated-like ensemble with weak CD density, and the preactivated-like ensemble with weak TMD density, we performed 3DVA with a full channel mask and filter resolution between 5-8 Å and clustered each of 3 modes independently into 5 groups. Occasionally, one or two clusters would be populated with very few particles, suggesting that a fewer number of clusters was adequate to represent the underlying heterogeneity. We then refined each class (CryoSparc v3.3.1 Local Refinement due to symmetry expansion) and assessed the resulting structures, selecting the mode of variability that contained our features of interest. From the resting-like and preactivated-like stacks with weak TMD density, we obtained the ∼C2 and ∼C4 TMD transition states presented in Figure 4D-E and S15. From the resting-like and preactivated-like stacks with weak CD density, we obtained the asymmetric ARM2 sampling states presented in Figure 3A. For the ARM2 retraction analysis in Figure 3B-G and wedge loop analysis in Figure S12K-P, we increased the requested number of clusters to 10 and 20 respectively and selected 6 refinements that appeared to be on a shared trajectory for both ARM2 retraction and loop melting for presentation.

For resolving the higher-order interactions between inhibited particles, the non-symmetry expanded stack of 1045k particles was subjected to CryoSparc v3.3.1 Heterogeneous Refinement seeded with 24 identical references of the inhibited state, which resulted in three classes (246k particles total) with strong density for interactions with an adjacent tetramer. This classification was used to quantify the particle distributions for clustered versus isolated inhibited states. The clustered particles were subjected to C4 symmetry expansion and local refinement using the entire CD mask, which showed that the interaction was likely formed between ARM2 of the central protomer and CLD of the adjacent protomer. Creating a new soft mask of these two interacting domains, we performed local refinement, 3DVA, and clustering along the tertiary mode of variability into 5 groups to resolve the ordered interaction at 3.4 Å from 79k particles. We then applied density modification and built a model of the interaction from two tetramers spanning residues 790-1696 from the central tetramer and residues 644-1695 from the adjacent tetramer.

For the depictions of the composite maps in Figure 1, four copies of the single-chain composite were fit to the consensus C4 refinement and combined using the Chimera VOPMAX command ^107^. For the depictions of ARM2 density in Figure 3, unsharpened local CD refinements were shown. For depictions of the overall density or slices at the JD ring, gate, and S1’-S1’’ in Figure 4 and Figure S15, the unsharpened consensus refinement maps were low-pass filtered to 4 Å (overall) or 7 Å (zoomed) using relion_image_handler ^109, 110^. For the depictions of the wedge loop density in Figure S13B, the resting state composite map was used. For the depictions of the wedge loop density in Figure S13K-P, a B-factor derived from the Guinier plot was used to sharpen the CD local refinements for presentation. All figures depicting models were generated in PyMol (Schrodinger, LLC. 2010. The PyMOL Molecular Graphics System, Version 2.5.3), and all figures depicting density alone were generated in ChimeraX ^111, 112^. For the electrostatics calculations in Figure S14C-D and S17A, the Adaptive Poisson-Boltzmann Solver (APBS) algorithm ^113^ was utilized via PyMol plug-in.

### High performance computing

The MSK HPC resource provides a GPU cluster built for computing large volume data over a range of applications from drug discovery to deep learning and image processing. It contains 120 nodes connected by a 100 Gigabit ethernet backbone. The nodes used for this project each contain Intel Xeon Platinum 2.2 GHz CPUs and 1 TB DDR4 RAM. Each node also contains four A100 GPUs interconnected using NVLink. The cluster runs the CentOS operating system and is supported by a 4 PB high-speed GPFS-based parallel filesystem. A 200 TB NVMe-based Weka ultra-fast tier was used as scratch space. The CPU to GPU communication is established over PCIE 4.0. The project used IBM Spectrum LSF as the orchestrator of shared resources and parallelization is further achieved by MPI over the ethernet network. All cryo-EM software excluding CryoSparc was maintained via HMS SBGrid^114^. Multiple sequence alignments were performed using the MUSCLE ^115^ algorithm in DNASTAR LaserGene MegAlign Pro 17.3.

### Adherent cell culture

HEK293T-IP_3_R-null cells were obtained through Kerafast ^51^ and cultured to a confluency of ∼75-80% on 100 × 20 mm tissue culture treated dishes in DMEM supplemented with 10% fetal bovine serum, 100 U/ml penicillin, 100 mg/ml streptomycin at 37°C with 5% CO_2_. For imaging, cells were then split in a 1:4 ratio and plated on poly-D-lysine coated, 35mm diameter, optical quality glass-bottom culture dishes (World Precision Instruments; #FD35PDL-100) and incubated for ∼18-24 hours. At ∼60% confluency, cells were transduced with a 200 µl baculovirus followed by incubation at 37 C, 5% CO_2_ for another 24 hours. All constructs used for Ca^2+^ imaging in this study were overexpressed in HEK293T-IP3R-null cells using the BacMam system ^116^.

### Ca^2+^ imaging and data processing

24 hours after baculovirus transduction, cells were gently washed with imaging buffer [20 mM HEPES supplemented Ca^2+^, Mg^2+^ free, Hank’s balanced salt solution (ThermoFisher; #14175103)] followed by incubation for 1 hour at 37°C and 5% CO2 in 1800 µl of imaging buffer containing 3 mM Cal-520-AM (AAT Bioquest; #21130) Cal-520-AM-loaded cells were removed from the CO2 incubator and equilibrated at room temperature for 5 minutes prior to IP_3_ stimulation by the addition of 200 µl of 1 mM carbachol (Alfa Aesar; #L06674-06), a Gαq-coupled M3 muscarinic receptor agonist. Carbachol was added at least 10 mm away from the imaging site and allowed to diffuse to a final concentration of 100 µM. Movies of carbachol-induced Ca^2+^ release in cells were collected at 20x with LD Plan-Neofluar 20X/0.4 Korr M27 objective, for 10 minutes, at 3×3 binning (912×736 pixels post binning), with an exposure time of 250ms on a Zeiss Axio observer D1 inverted phase-contrast fluorescence microscope equipped with an Axiocam 506 Mono camera (Zeiss). Cal-520-AM imaging was carried out by exciting the sample at 493 nm and monitoring emission at 515 nm using X-Cite Series 120Q illumination system and Zeiss filter set 38 HE.

Ca^2+^ imaging movies were processed using ImageJ ^117^, Fiji ^118^ and MathWorks MATLAB 9.12.0.1884302 (R2022a) to extract Cal-520-AM fluorescence traces from individual cells. Movie stacks were background-subtracted with a 200-pixel rolling ball radius in ImageJ. Maximum intensity projection of the stack was used to generate a difference of gaussian image, which was used for edge detection and cell segmentation using MATLAB’s Image Processing Toolbox. Traces were then extracted from segmented cells, smoothed over 41 frames using a Savitzky–Golay filter of polynomial order 2, normalized by Z-score, and baseline adjusted using the linear method of MATLAB’s 1-D data interpolation function with a custom MATLAB script called Baseline Fit ^119^. In the baseline-adjusted traces, the smallest observed Ca^2+^ oscillation peak value was used to manually threshold and identify other peaks automatically. Detected peaks were then used to calculate inter-spike intervals using MATLAB’s Signal Processing Toolbox. All statistical tests were performed using GraphPad Prism 9. Data reported are from 3 independent biological replicates.

For analysis of peaks from individual replicates, traces with transients/oscillations were baseline adjusted in MATLAB using Baseline Fit ^119^ and normalized between 0 and 1. The first peak of each oscillation/transient was identified and a window of 75 seconds on both sides of the peak was extracted and aligned at the peak position. Mean and 95% confidence intervals were calculated using GraphPad Prism and overlayed on traces from a single biological replicate. A 1 second window on both sides of the mean data point corresponding to half maximal intensity were fit to a straight line and used to calculate the mean rising phase for constructs exhibiting transients/oscillations. Traces with oscillations were sorted based on the maximum number of distinguishable peaks and plotted as a fraction of total oscillatory traces.

### Fura-2 calibration

10 µl of the 5X ligand and Ca^2+^ chelator cocktail (described earlier) of 100 – 4 nM free Ca^2+^ concentration was added to 40 µl of 62.5 µM Fura2 (ThermoFisher; #F-1200) diluted in gel filtration buffer [150 mM NaCl, 50 mM Tris-HCl pH 8.0, 0.02% LMNG, and 2 mM dithiothreitol (DTT)] to bring Fura-2 to a final concentration of 50 mM. Samples were excited at 340 nm and 380 nm, and fluorescence emissions were collected at 510 nm in a 96 well black/clear bottom plate using Molecular Devices SpectraMax M5e microplate reader at room temperature. Fluorescence emission ratios at 340/380 excitation were then calculated and fitted to a sigmoid using a logistic dose-response function in GraphPad Prism 9.

