## Supplementary Figures and Tables for "Structural titration reveals Ca^2+^-dependent conformational landscape of the IP_3_ receptor"

#### ***Supplemental Figures:***

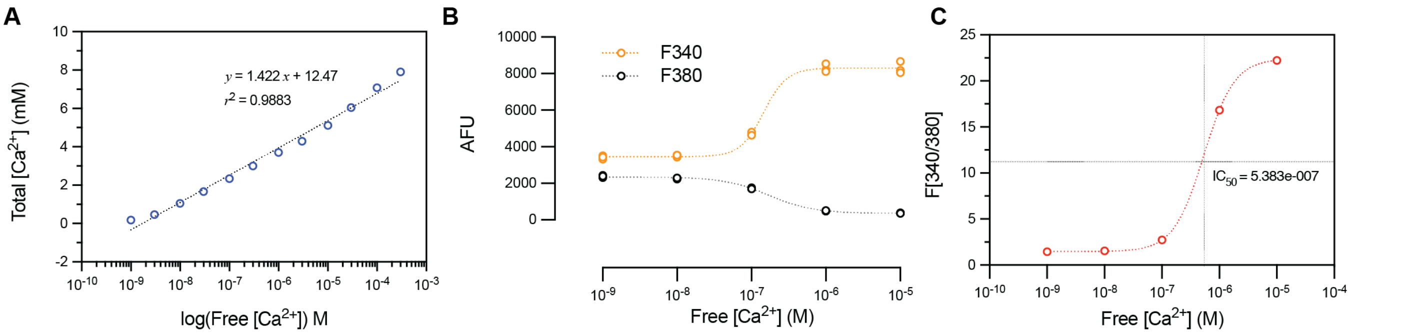

**Figure S1: Modeling and experimental assessment of the $Ca^{2+}$ buffering system. (A)** Modeling of free versus total $Ca^{2+}$ in the experimental buffer using MaxChelator. Points were fit to a line with $r^2$=0.99. **(B)** Fura-2 measurements of free $Ca^{2+}$ concentrations in the experimental buffer. Plot shows Fura-2 510 nm emissions for each $Ca^{2+}$ condition excited at 340 nm and 380 nm. **(C)** Semi-log plot of Fura-2 340/380 nm ratios versus expected free $Ca^{2+}$ concentration.

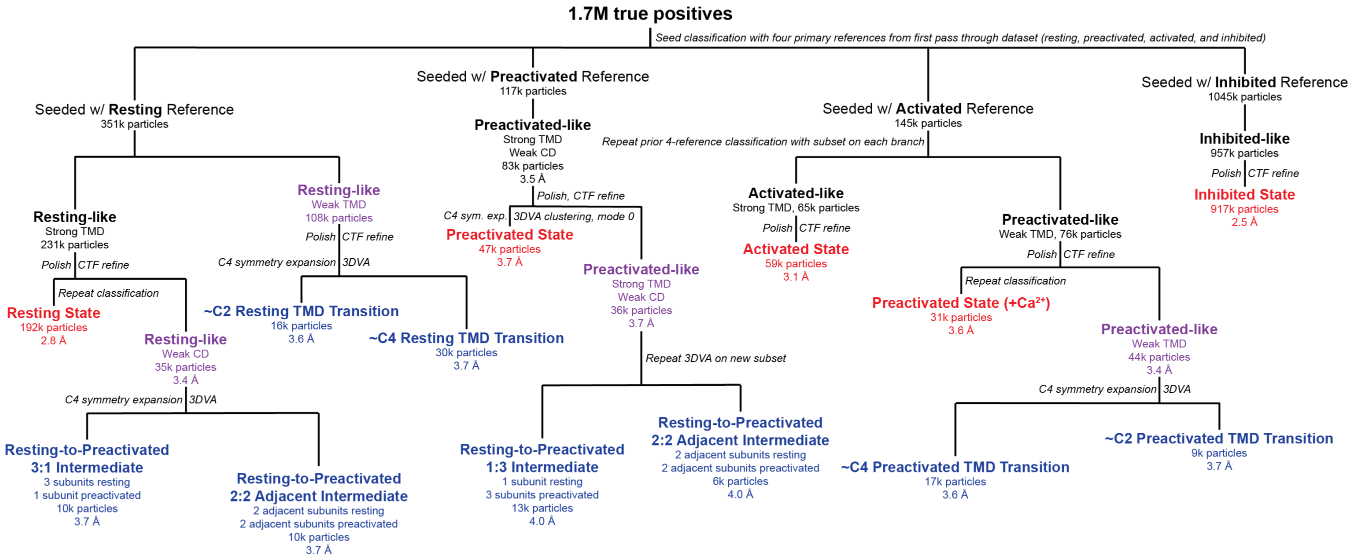

**Figure S2: Cryo-EM classification workflow highlighting key observations and decision-making criteria.** When appropriate, particle numbers are presented along with $FSC_{0.143}$ refinement resolution. States depicted in red are the primary C4 symmetric states. Classes depicted in purple are conformational ensembles that were investigated using 3DVA. States depicted in blue are conformational states obtained from 3DVA that are derived from the preceding ensemble.

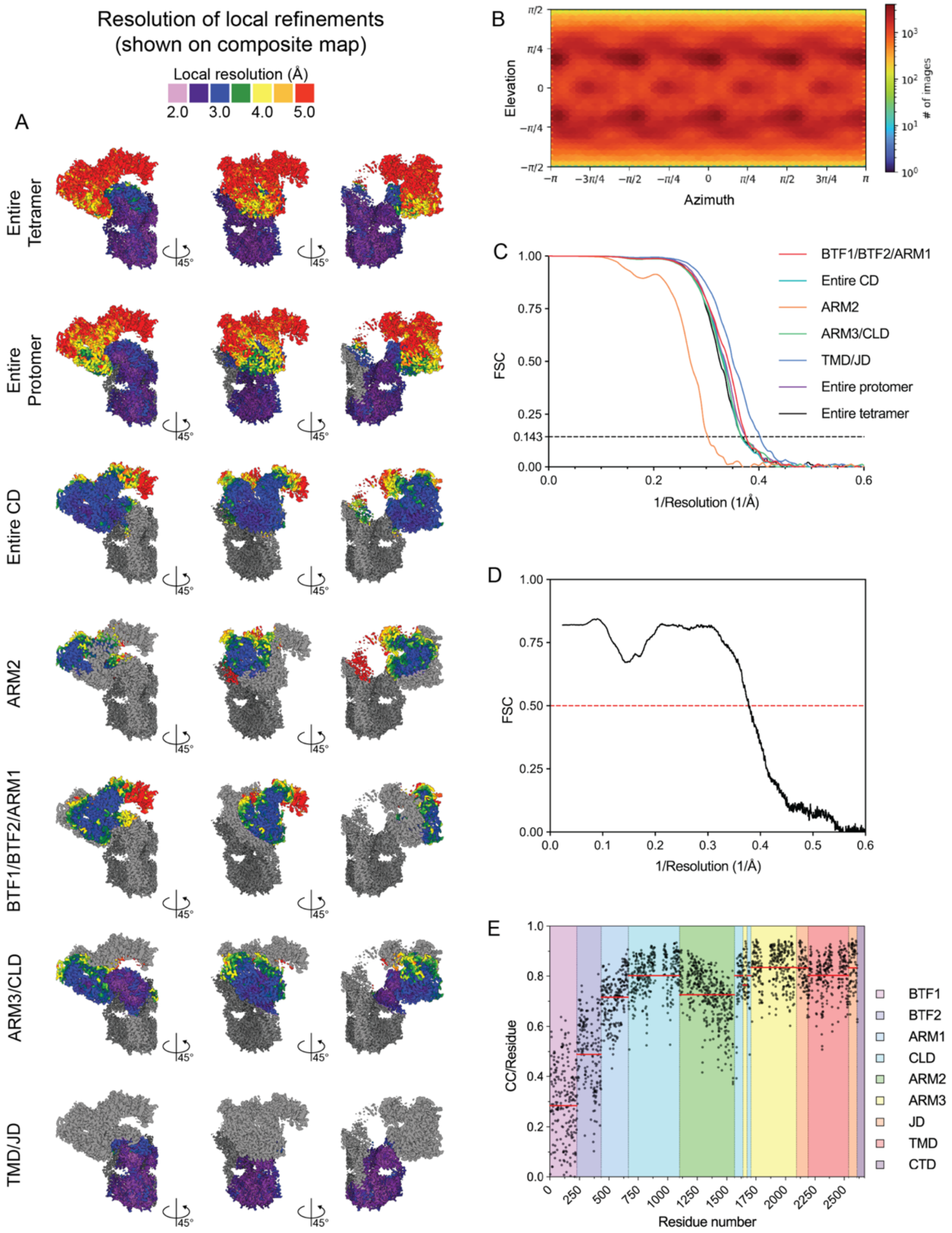

**Figure S3: Validation of the inhibited state. (A)** Local resolution plots at $FSC_{0.5}$ for specific local consensus or local refinements depicted on the final density modified composite map. **(B)** Angular distribution plot for the consensus refinement. **(C)** Half map $FSC_{0.143}$ plots for the local refinements. **(D)** $FSC_{0.5}$ map versus model. **(E)** Per-residue cross correlation (CC) for map-to-model comparison with domain demarcations and a red bar highlighting the average CC for each domain.

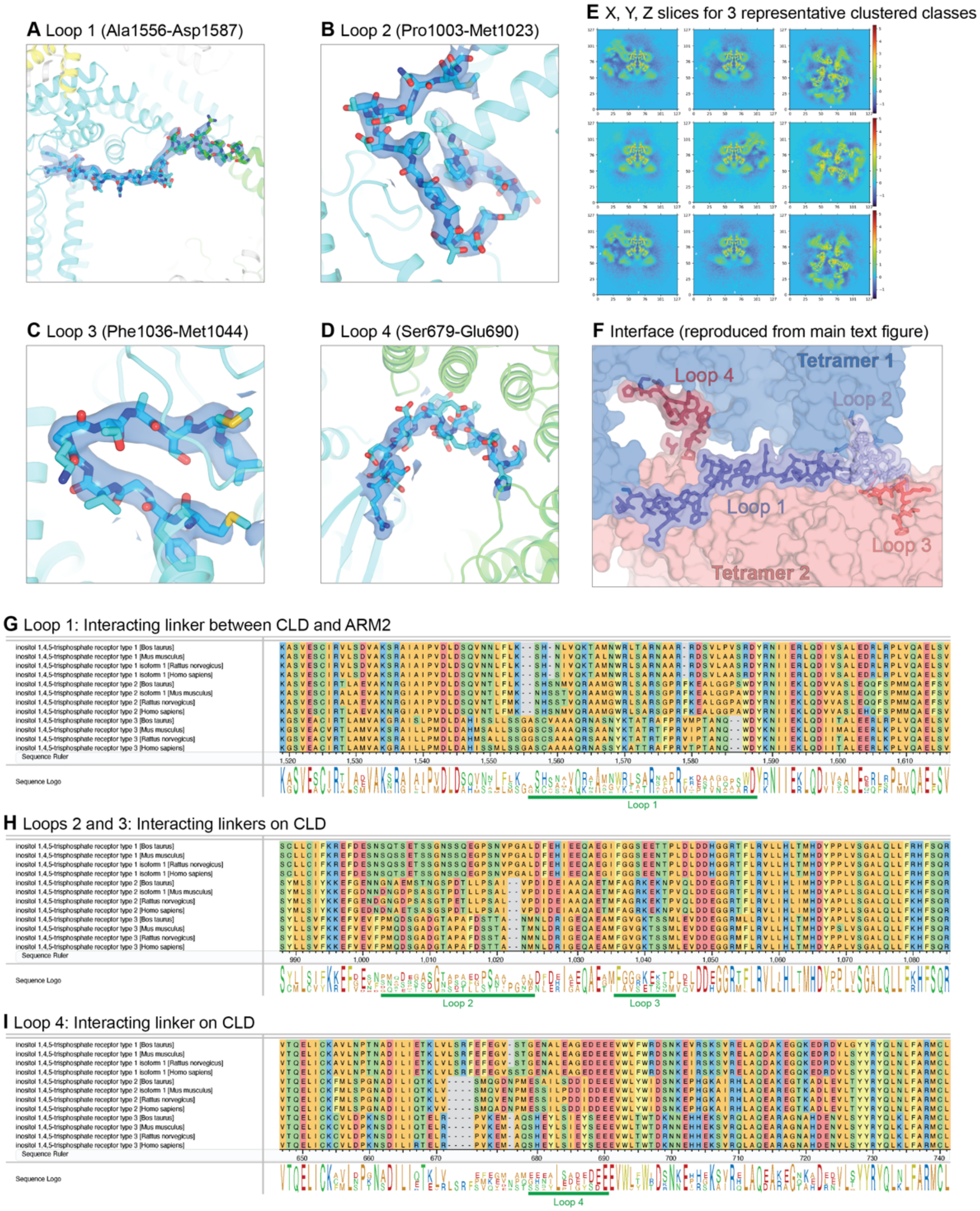

**Figure S4: Higher-order interactions between inhibited particles form clusters. (A-D)** Structures of (A) loop 1, (B) loop 2, (C) loop 3 and (D) loop 4 depicted as sticks. Cryo-EM density shown as blue isosurfaces **(E)** Central slices through the density for representative classes of the clustered inhibited state showing strong density for an interaction with an adjacent protomer that extends outside of the refinement box. **(F)** Model of the interacting loops between two tetramers (colored pink and blue) with the loops contributed from each shown as a similar color. **(G)** Multiple sequence alignment for region between ARM2 and the CLD that contributes loop 1. **(H)** Multiple sequence alignment for the region of the CLD contributing both loop 2 and loop 3. **(I)** Multiple sequence alignment for the region of the CLD contributing loop 4.

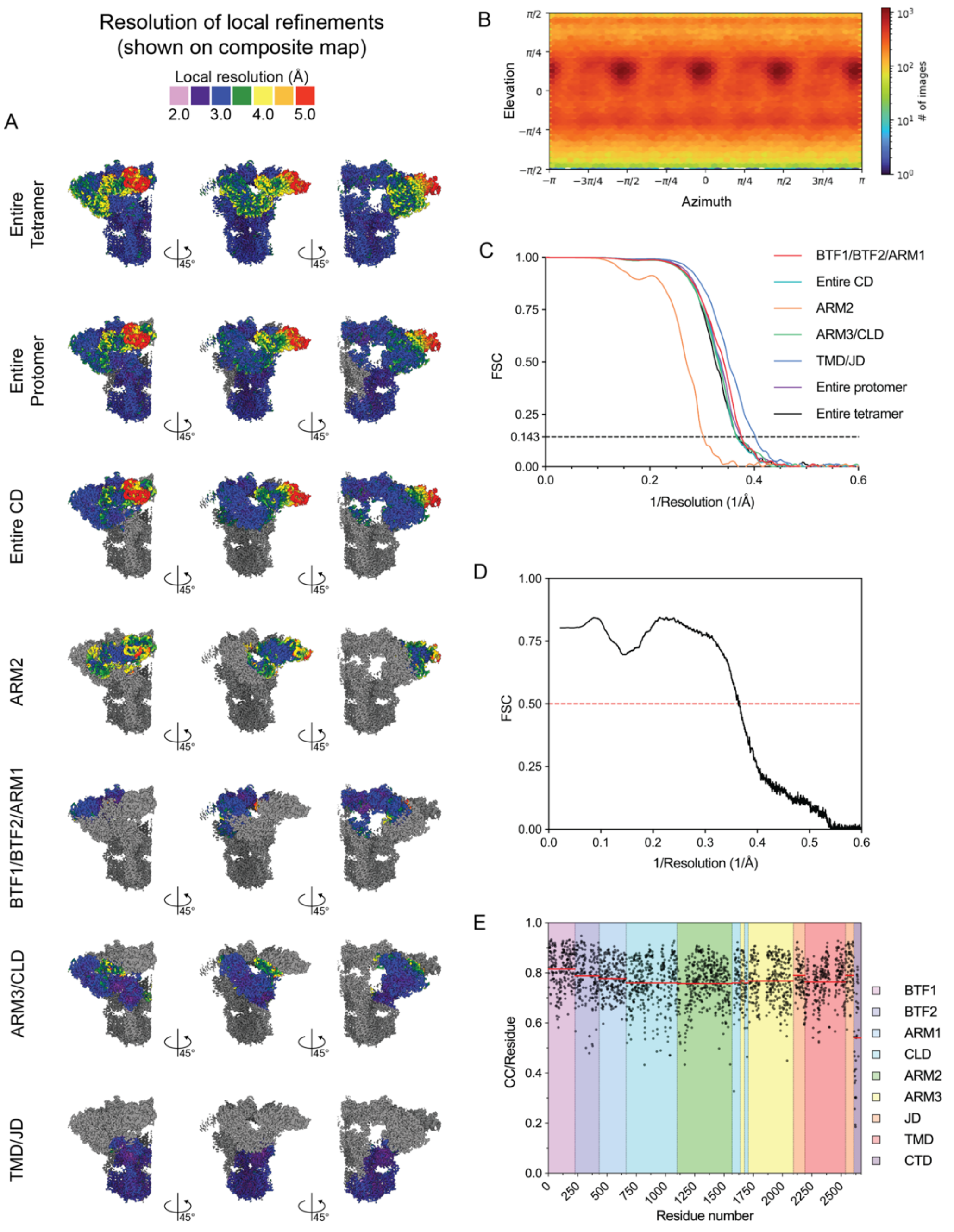

**Figure S5: Validation of the resting state. (A)** Local resolution plots at $FSC_{0.5}$ for specific local consensus or local refinements depicted on the final density modified composite map. **(B)** Angular distribution plot for the consensus refinement. **(C)** Half map $FSC_{0.143}$ plots for the local refinements. **(D)** $FSC_{0.5}$ map versus model. **(E)** Per-residue cross correlation (CC) for map-to-model comparison with domain demarcations and a red bar highlighting the average CC for each domain.

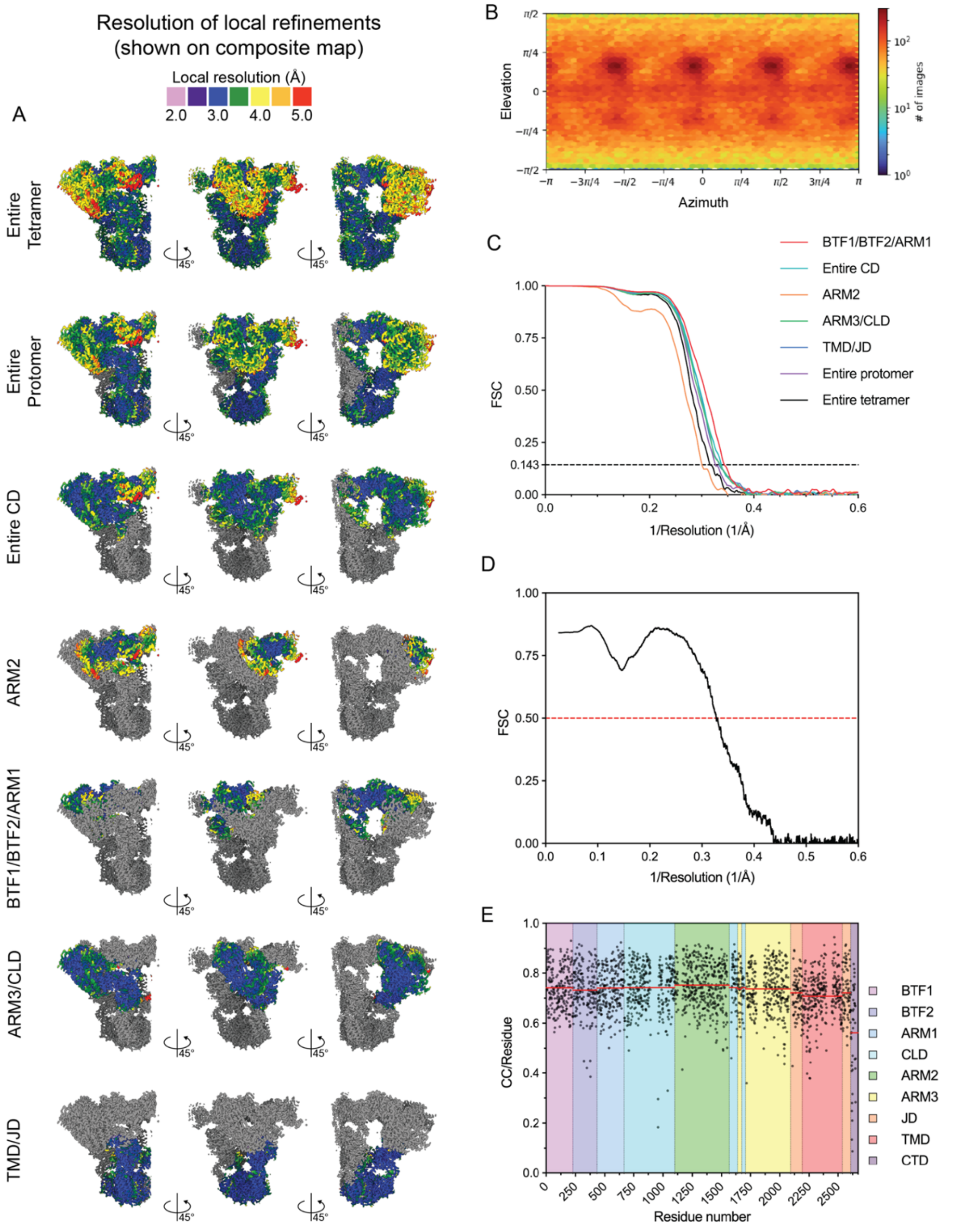

**Figure S6: Validation of the activated state. (A)** Local resolution plots at $FSC_{0.5}$ for specific local consensus or local refinements depicted on the final density modified composite map. **(B)** Angular distribution plot for the consensus refinement. **(C)** Half map $FSC_{0.143}$ plots for the local refinements. **(D)** $FSC_{0.5}$ map versus model. **(E)** Per-residue cross correlation (CC) for map-to-model comparison with domain demarcations and a red bar highlighting the average CC for each domain.

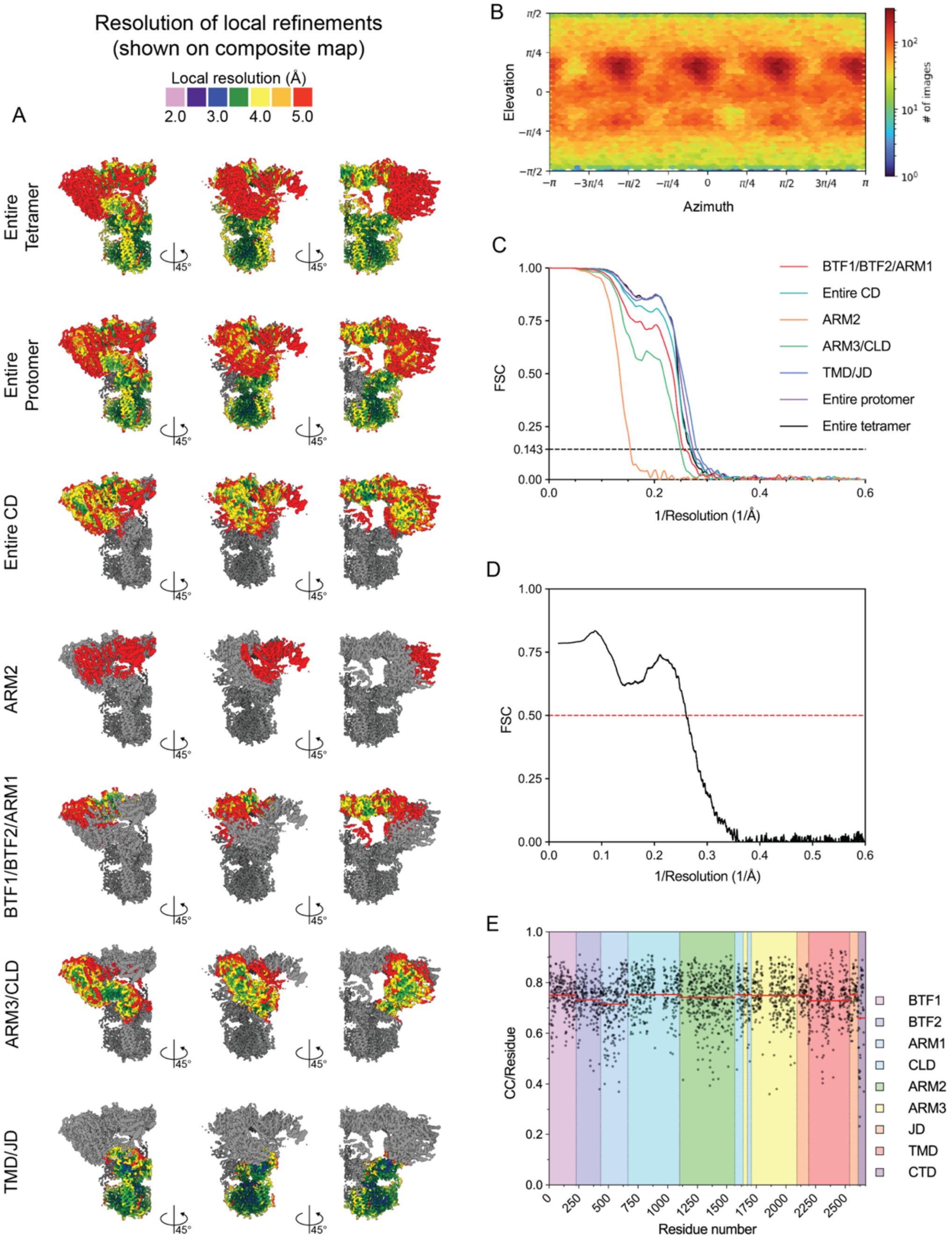

**Figure S7: Validation of the preactivated state. (A)** Local resolution plots at $FSC_{0.5}$ for specific local consensus or local refinements depicted on the final density modified composite map. **(B)** Angular distribution plot for the consensus refinement. **(C)** Half map $FSC_{0.143}$ plots for the local refinements. **(D)** $FSC_{0.5}$ map versus model. **(E)** Per-residue cross correlation (CC) for map-to-model comparison with domain demarcations and a red bar highlighting the average CC for each domain.

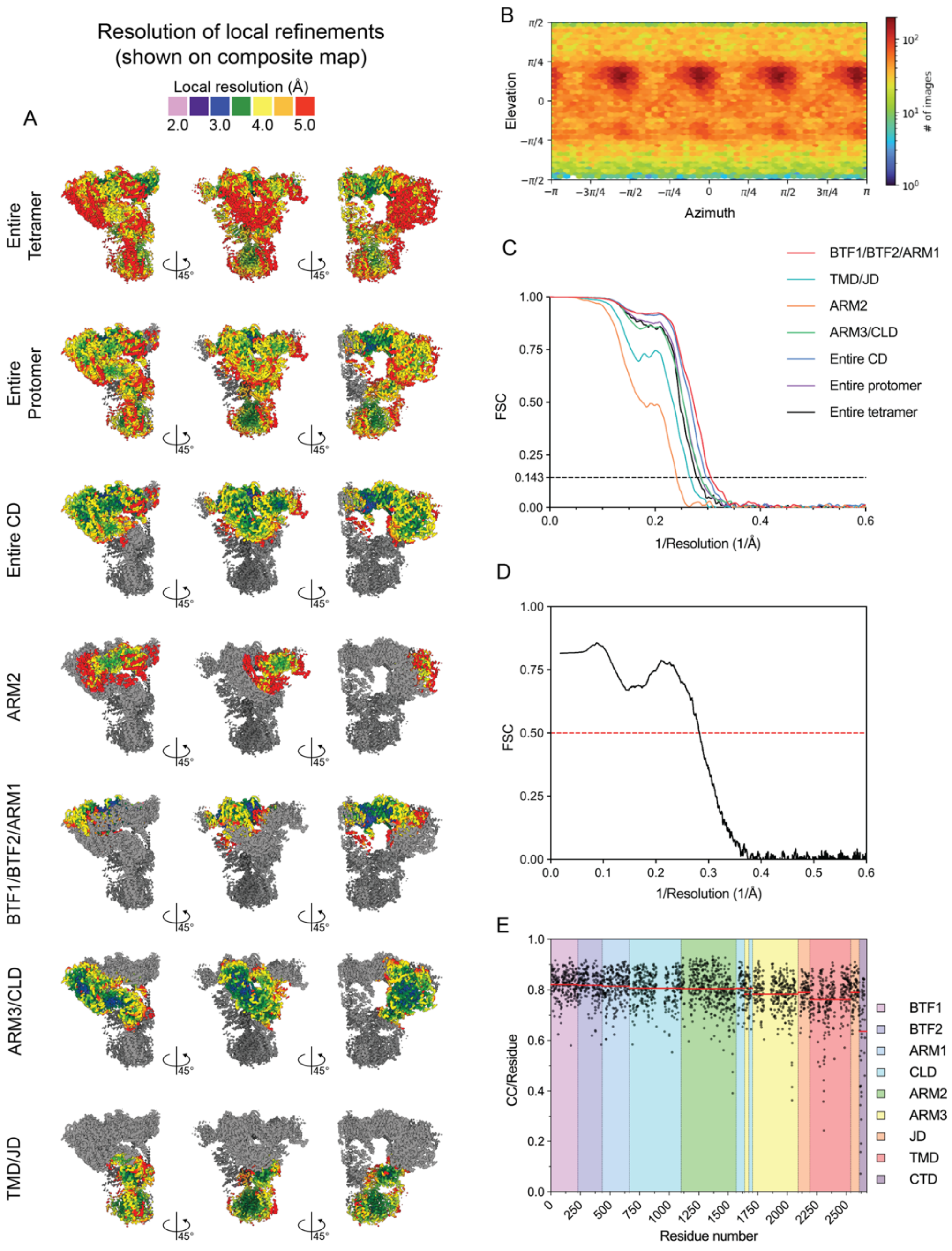

**Figure S8: Validation of the preactivated+Ca$^{2+}$ state. (A)** Local resolution plots at $FSC_{0.5}$ for specific local consensus or local refinements depicted on the final density modified composite map. **(B)** Angular distribution plot for the consensus refinement. **(C)** Half map $FSC_{0.143}$ plots for the local refinements. **(D)** $FSC_{0.5}$ map versus model. **(E)** Per-residue cross correlation (CC) for map-to-model comparison with domain demarcations and a red bar highlighting the average CC for each domain.

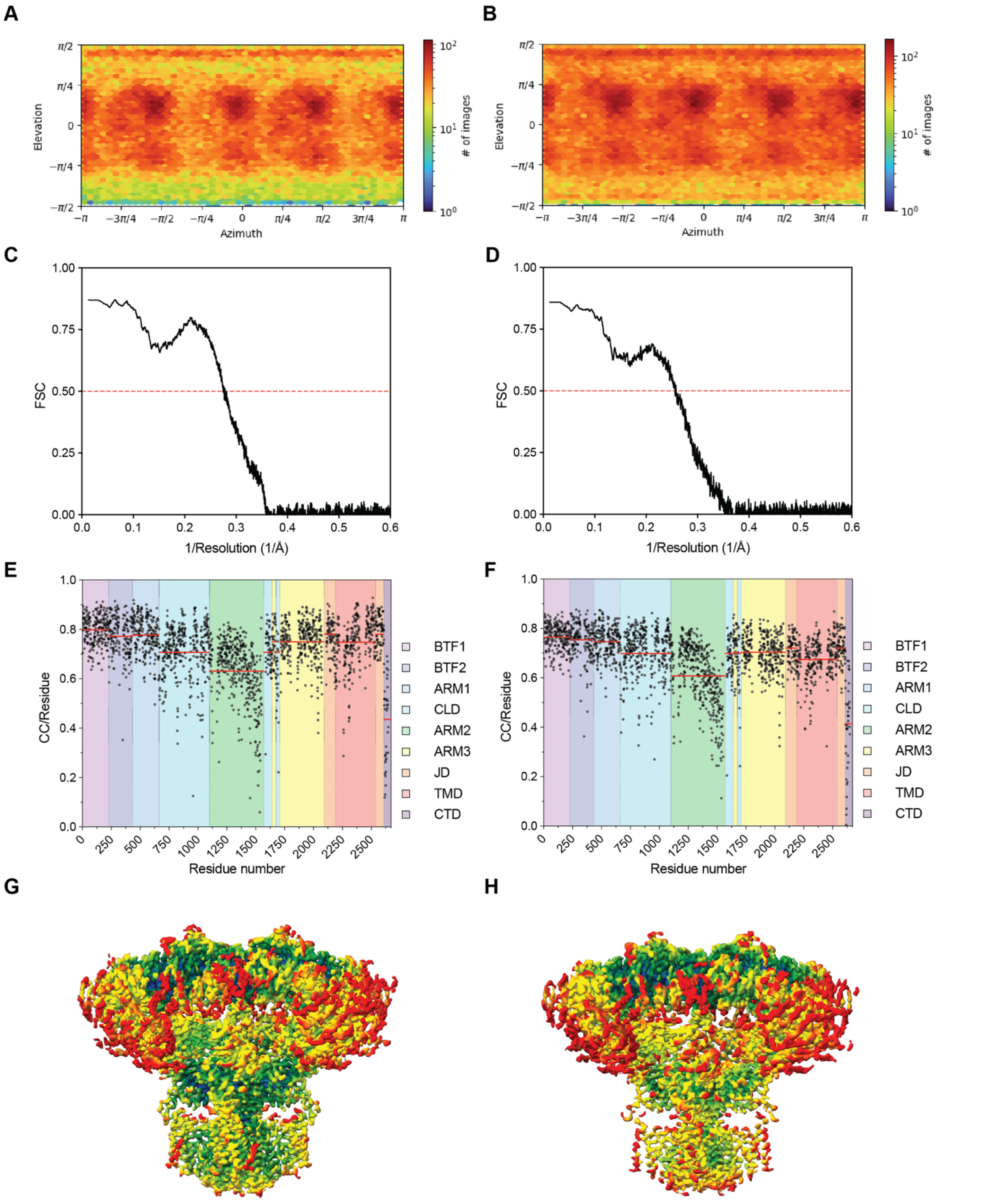

**Figure S9: Validation of the labile resting states.** (**A-B)** Angular distribution plot for the consensus refinement of (A) labile resting state #1 and (B) labile resting state #2. **(C-D)** $FSC_{0.5}$ map versus model for (C) labile resting state #1 and (D) labile resting state #2. **(E-F)** Per-residue cross correlation (CC) for map-to-model comparison with domain demarcations and a red bar highlighting the average

CC for each domain for (E) labile resting state #1 and of (F) labile resting state #2. **(G-H)** Local resolution plots at $FSC_{0.5}$ for specific consensus refinements depicted on the final density modified composite map of (G) labile resting state #1 and (H) labile resting state #2.

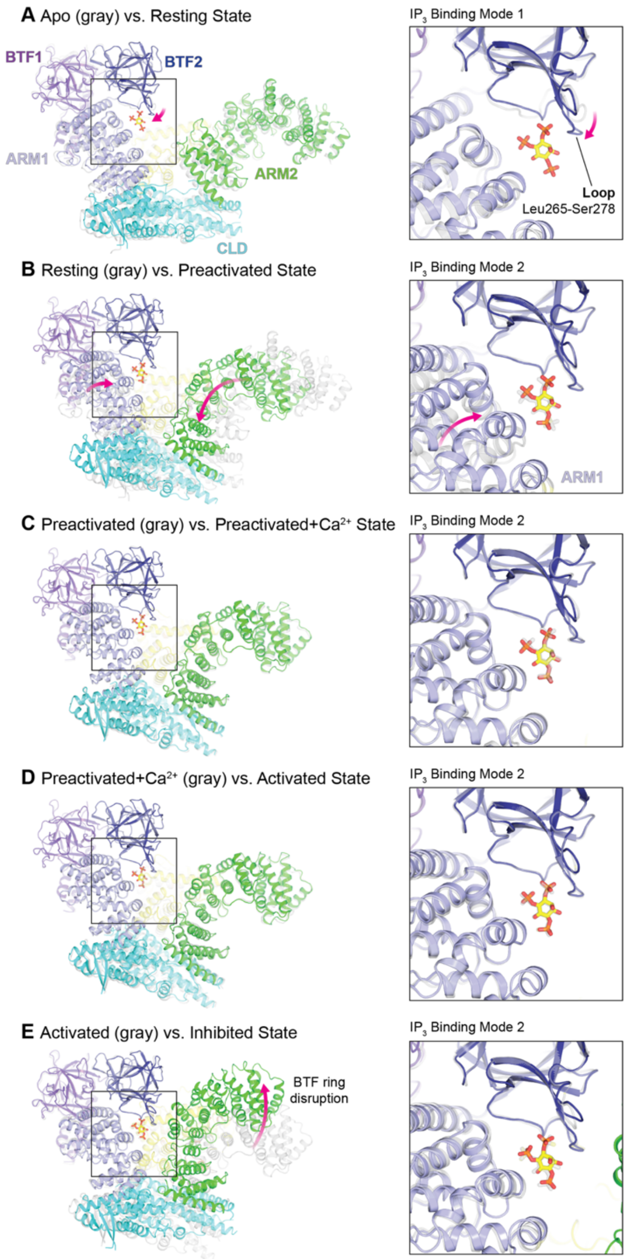

**Figure S10: $IP_3$ is coordinated in the binding pocket in two modes.** Overlay of the entire CD **(left)** and zoomed on the $IP_3$ binding pocket **(right)** for the comparisons between **(A)** resting and preactivated states **(B)** preactivated and preactivated+$Ca^{2+}$ states **(C)** preactivated+$Ca^{2+}$ and activated states **(D)** activated and inhibited states.

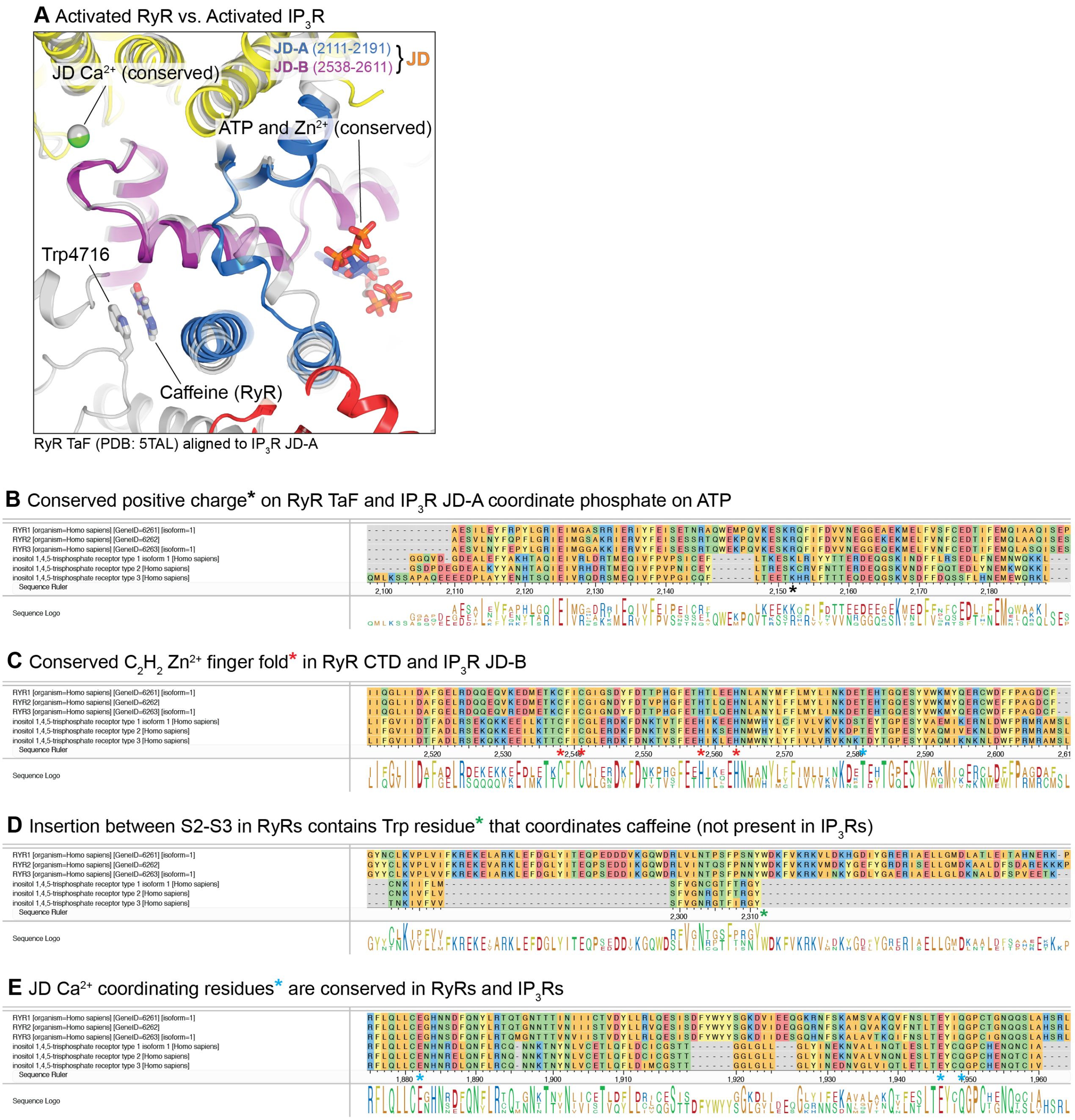

**Figure S11: Comparisons between the JD ligand binding sites in IP₃Rs and RyRs. (A)** Overlay of rabbit RyR1 (gray) and hIP$_3$R3 (colored) in their activated states highlighting the conserved JD $Ca^{2+}$, ATP, and $Zn^{2+}$ binding sites. The RyR caffeine binding site is not observed in hIP$_3$R3. The JD, composed of JD-A (blue) and JD-B (purple) are homologous to the thumb-and-forefinger (TaF) and C-terminal domains (CTD) in RyRs. The models were aligned by the TaF and JD-A. **(B-E)** Multiple sequence alignments for IP$_3$Rs and RyRs highlighting (B) conserved positive charge that coordinates phosphate on ATP (black asterisk), (C) conserved $C_2H_2$ $Zn^{2+}$ finger fold that coordinates $Zn^{2+}$ ion (red asterisks) (D), insertion in RyRs that forms caffeine binding site via a trytophan (green asterisk), and (E) conserved JD $Ca^{2+}$ coordinating residues (cyan asterisks).

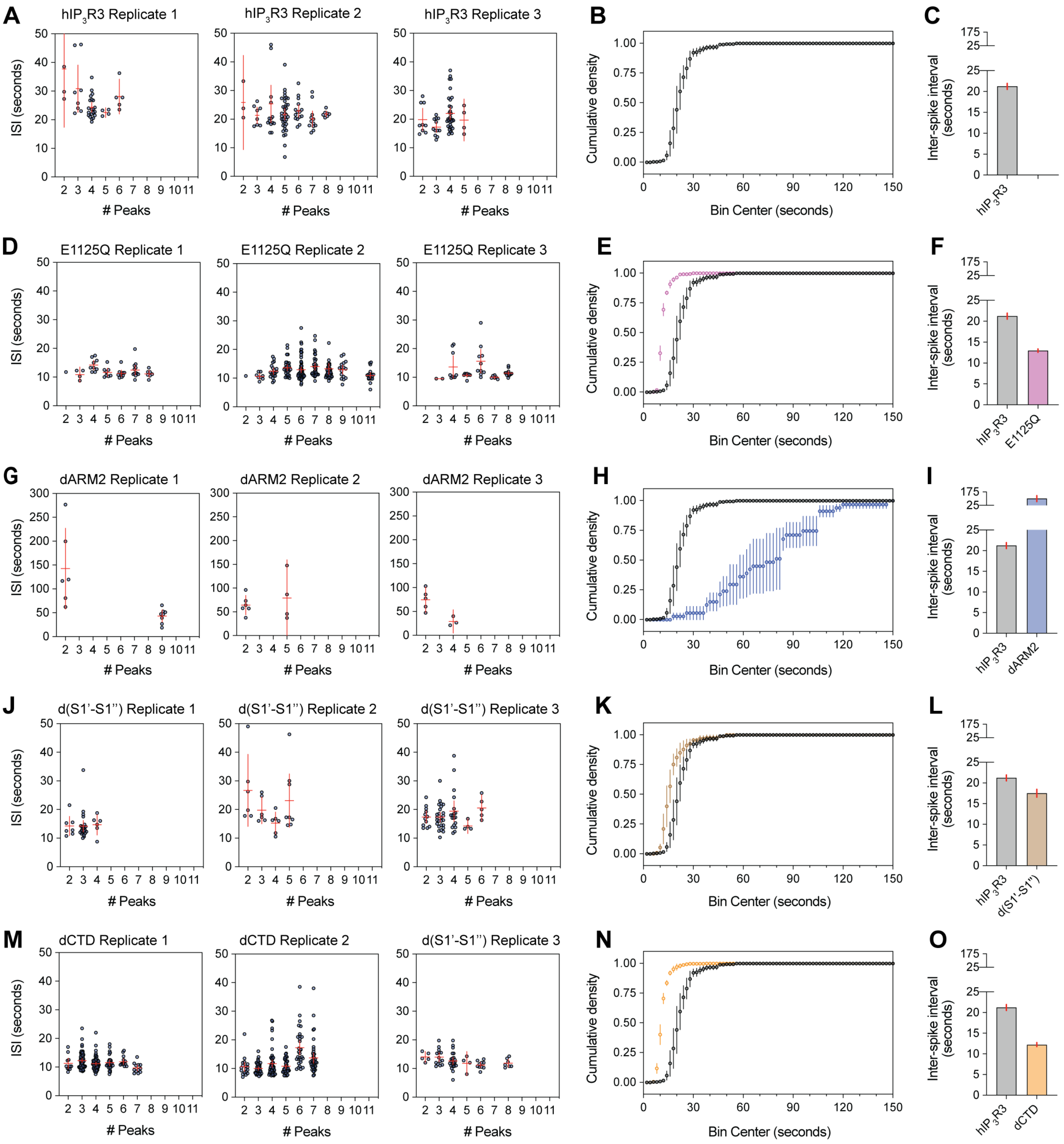
A
hIP3R3 Replicate 1
hIP3R3 Replicate 2
hIP3R3 Replicate 3
ISI (seconds)
### Peaks
B
Cumulative density
Bin Center (seconds)
C
Inter-spike interval (seconds)
hIP3R3
D
E1125Q Replicate 1
E1125Q Replicate 2
E1125Q Replicate 3
ISI (seconds)
### Peaks
E
Cumulative density
Bin Center (seconds)
F
Inter-spike interval (seconds)
hIP3R3
E1125Q
G
dARM2 Replicate 1
dARM2 Replicate 2
dARM2 Replicate 3
ISI (seconds)
### Peaks
H
Cumulative density
Bin Center (seconds)
I
Inter-spike interval (seconds)
hIP3R3
dARM2
J
d(S1'-S1'') Replicate 1
d(S1'-S1'') Replicate 2
d(S1'-S1'') Replicate 3
ISI (seconds)
### Peaks
K
Cumulative density
Bin Center (seconds)
L
Inter-spike interval (seconds)
hIP3R3
d(S1'-S1'')
M
dCTD Replicate 1
dCTD Replicate 2
d(S1'-S1'') Replicate 3
ISI (seconds)
### Peaks
N
Cumulative density
Bin Center (seconds)
O
Inter-spike interval (seconds)
hIP3R3
dCTD

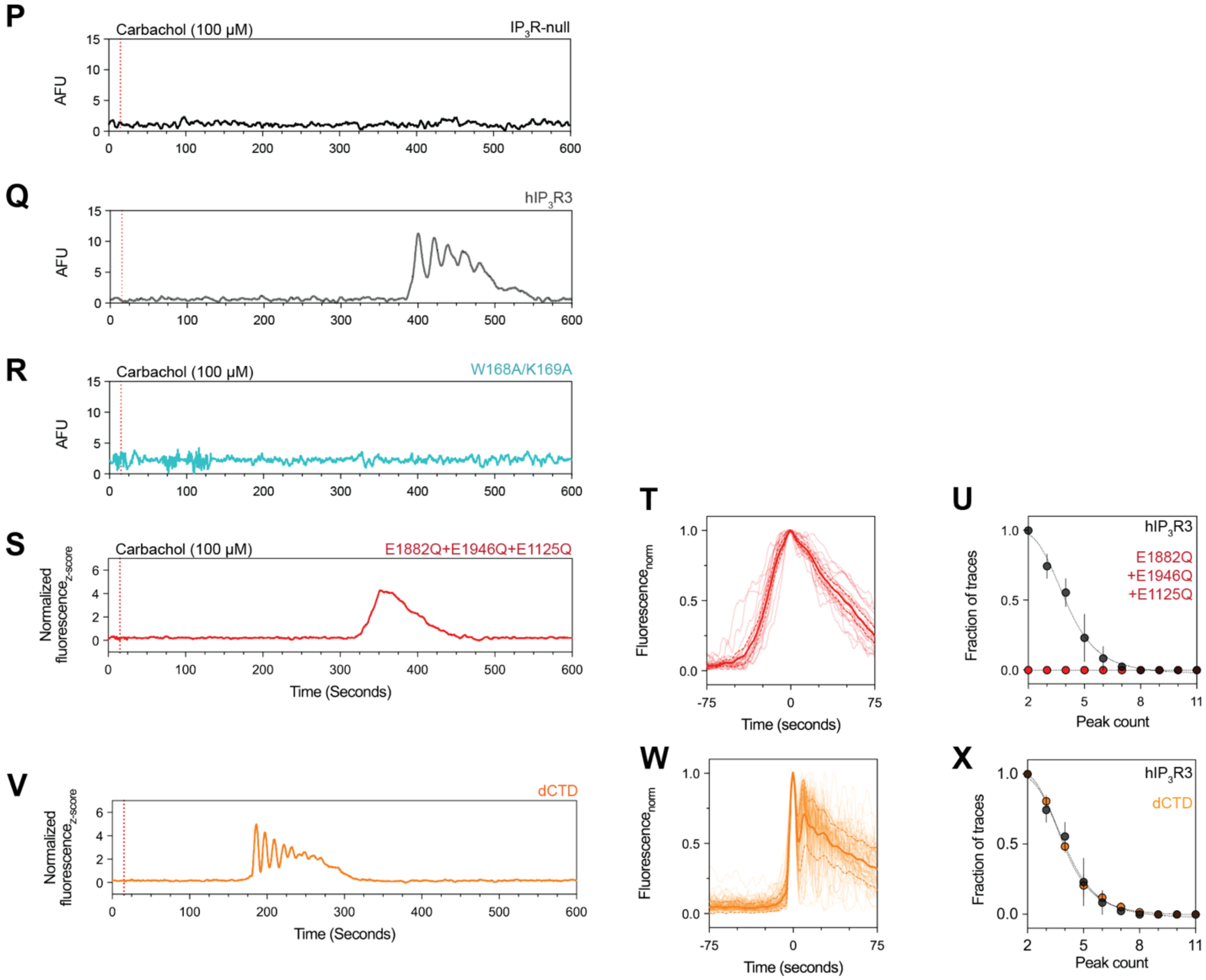

**Figure S12: $Ca^{2+}$ imaging data analysis. (A, D, G, J, M)** Whisker plot of inter-spike intervals for 3 independent biological replicates sorted by the minimum number of identifiable peaks observed in traces from which their values were obtained for (A) $hIP_3R3$, (D) E1125Q, (G) dARM2, (J) d(S1'-S1''), and (M) dCTD. Error bars correspond to 95% confidence interval around the mean. **(B, E, H, K, N)** Cumulative density plot for inter-spike intervals from 3 biological replicates of (B) hIP3R3, (E) E1125Q, (H) dARM2, (K) d(S1'-S1''), and (N) dCTD. Error bars correspond to 95% confidence interval around the mean. **(C, F, I, L, O)** Bar plots depicting the mean inter-spike interval values for (C) $hIP_3R3$, (F) E1125Q, (I) dARM2, (L) d(S1'-S1''), dCTD (O). Error bars correspond to 95% confidence interval around the S.E.M. **(P-R)** Representative filtered, and baseline adjusted traces for (P) $IP_3R$-null cell line, cells overexpressing (Q) $hIP_3R3$ and (R) W168A/K169A. **(S, V)** Representative z-score normalized Cal-520-AM fluorescence trace recorded from cells expressing (S) E1882Q+E1946Q+E1125Q and (V) dCTD mutant in an $IP_3R$-null background, following stimulation by carbachol**. (T, W)** Aligned first peak of every oscillatory trace normalized to 1 for (T) E1882Q+E1946Q+E1125Q and (W) dCTD. **(U,X)** Distribution of peak counts for all oscillatory traces for (U) E1882Q+E1946Q+E1125Q and (X) dCTD. Individual points represent mean and error bars represent S.E.M.

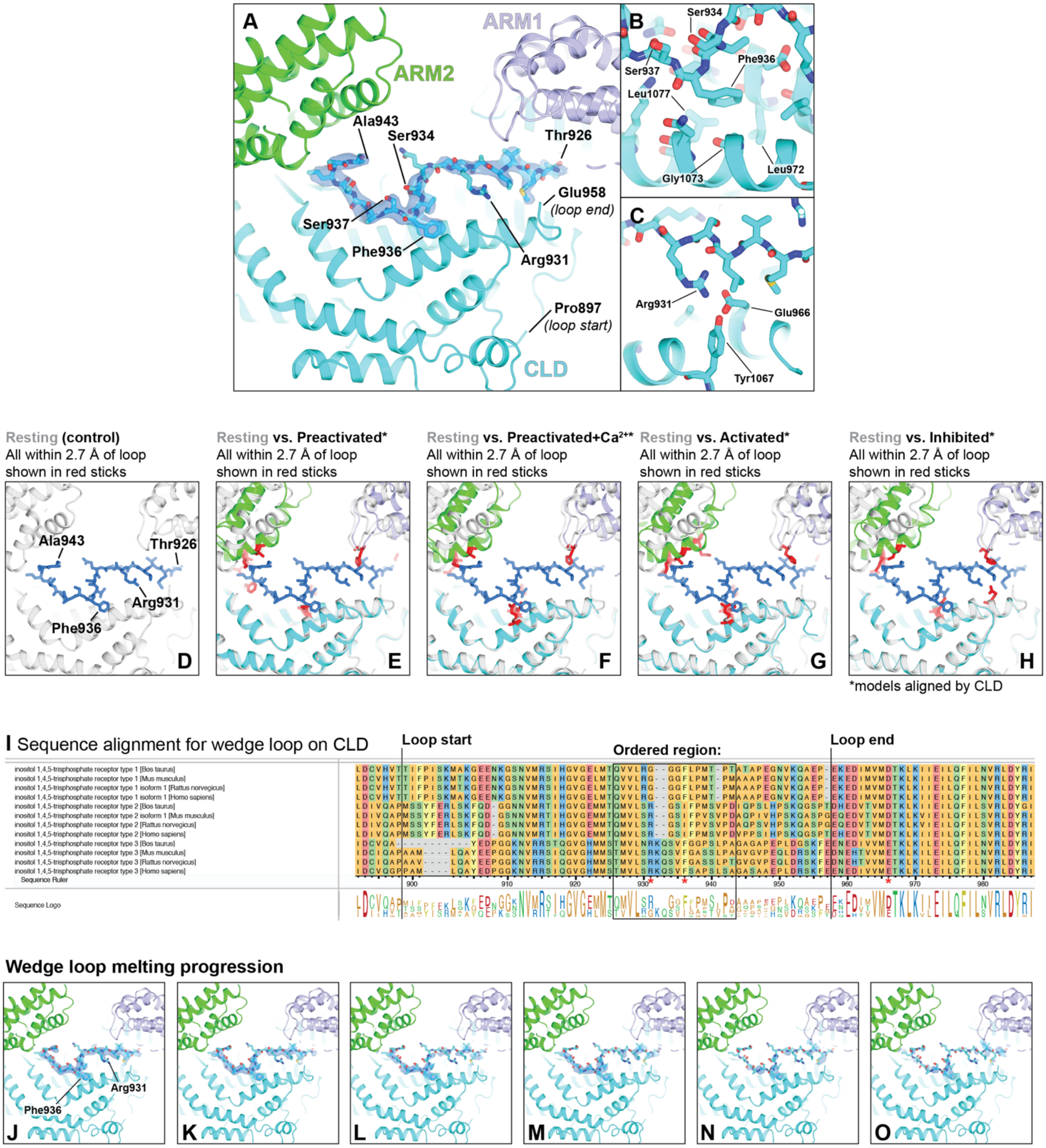

**Figure S13: Wedge loop prevents ARM2 retraction to stabilize the resting state. (A)** The wedge loop occupies a cavity between ARM1, ARM2 and the CLD in the resting state. Ordered residues within the wedge loop are depicted as sticks. Cryo-EM density for the wedge loop is shown as a blue isosurface. **(B-C)** Interactions that stabilize the wedge loop at (B) Phe936 and (C) Arg931. **(D-H)** Model of wedge loop docked from the resting state with alignment by the CLD. All residues with atoms closer than 2.7 Å are considered clashes and shown as red sticks for (D) resting, (E) preactivated, (F) preactivated+$Ca^{2+}$, (G) activated, and (H) inhibited states. **(I)** Multiple sequence alignment of the wedge loop. Arg931, Phe936 and Glu966 are highlighted with red asterisks **(J-O)** Expanded 3DVA trajectory of the ARM2 extended position shown as blue isosurfaces reveals progressive disordering of the wedge loop. **(J)** Most stable density for wedge loop. **(K-L)** Residues near Arg931 begin to weaken. **(M-N)** N- and C-termini of wedge loop begin to weaken. **(O)** Phe936 is the last remaining strong density feature.

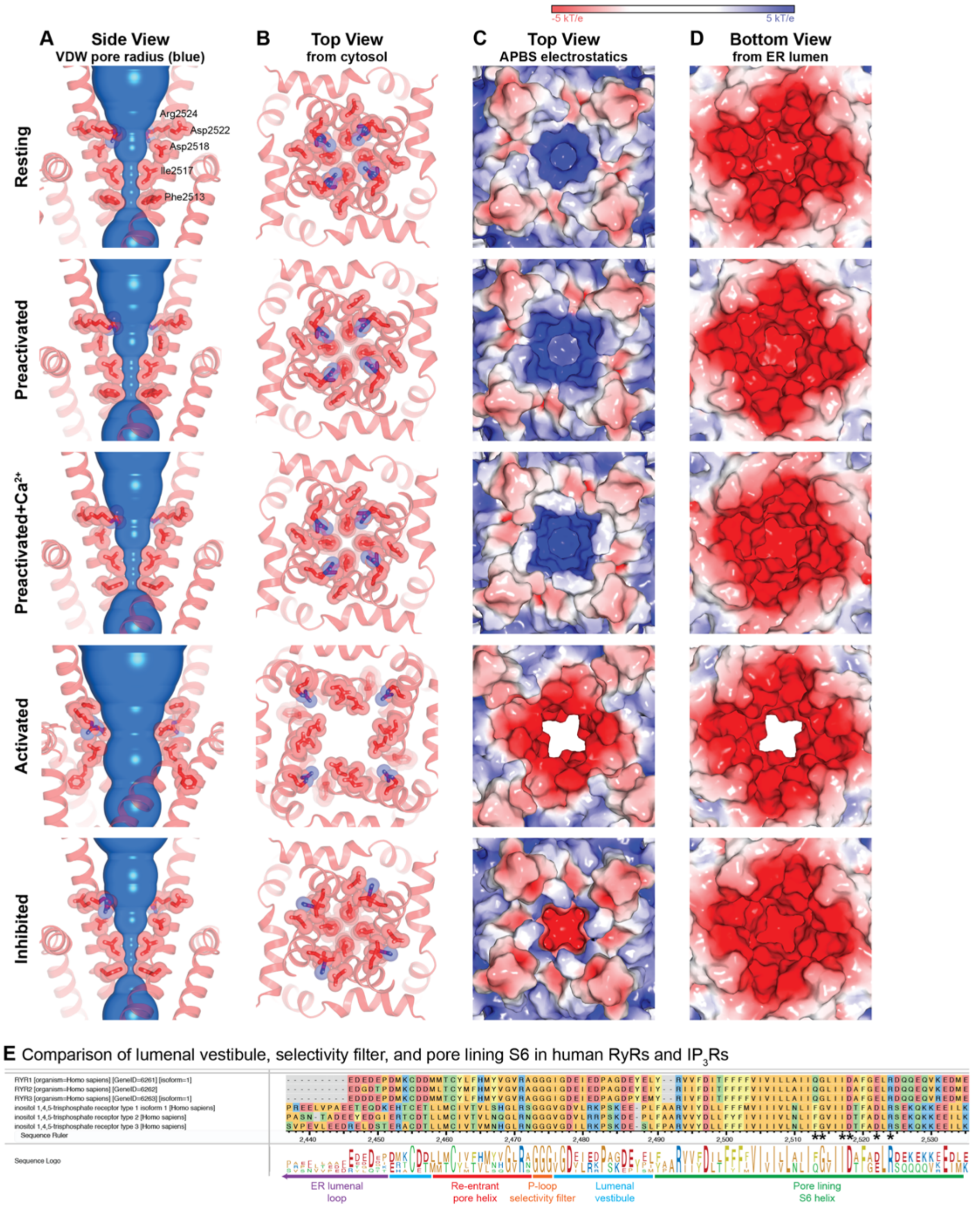

**Figure S14: Conformation and electrostatics of the pore. (A-B)** Structure of the pore viewed from the (A) side and (B) cytosol with front and rear protomers removed for clarity in A. Phe2513, Ile2517, Asp2518, Asp25222, Arg2524 shown as sticks and surfaces. **(C-D)** Surface depiction viewed from the (C) cytosol and (D) lumen colored by electrostatic surface potential in the cytosolic portion of the pore. **(E)** Multiple sequence alignment comparing luminal vestibule, pore helix, selectivity filter, and S6 helices in human $IP_3Rs$ and RyRs.

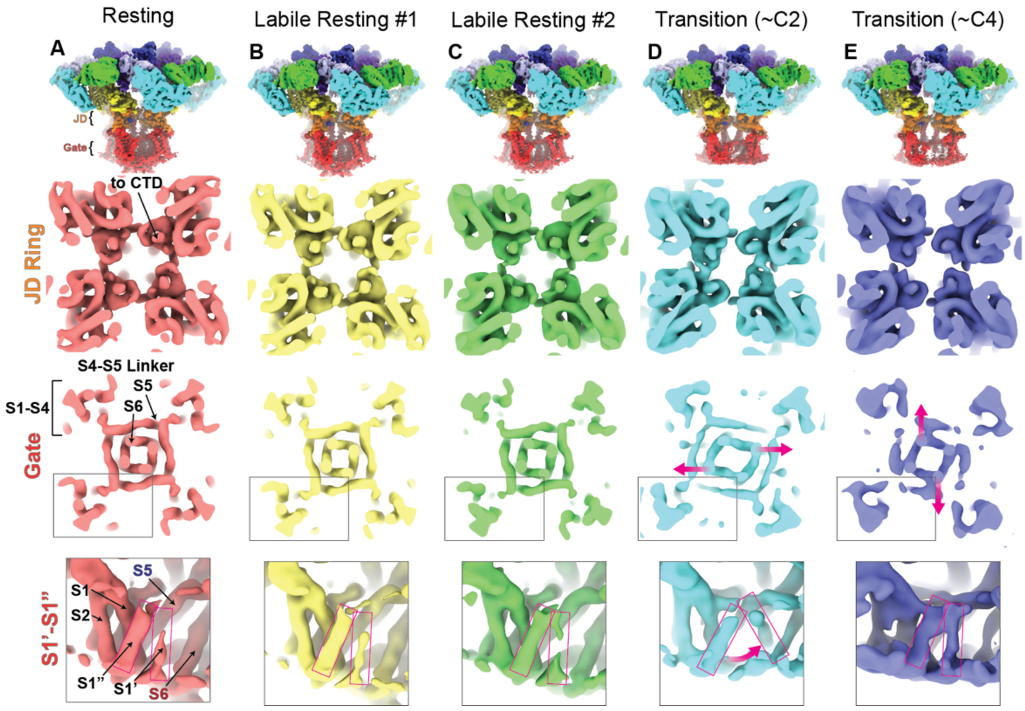

**Figure S15: Snapshots of conformational heterogeneity in the JD ring intact resting-like states. (A-F)** Cryo-EM density maps of the **(A)** resting state, **(B)** labile resting state #1, **(C)** labile resting state #2, **(D)** ~C2 resting TMD transition, **(E)** ~C4 resting TMD transition states, low-pass filtered to 4 Å (overall) or 7 Å (slices). **Row 1:** Overall cryo-EM density. **Row 2:** density slice looking from the cytosol at the height of the JD ring. **Row 3:** density slice looking from the cytosol at the height of the gate. **Row 4:** Side view of a single S1-S4 domain highlighting the position of S1'-S1" and the S4-S5 linker for the central protomer, with S5 from the adjacent protomer and S6 of the opposite protomer visible, highlighting the intertwined TMD of domain-swapped 6TM cation channels.

**A**

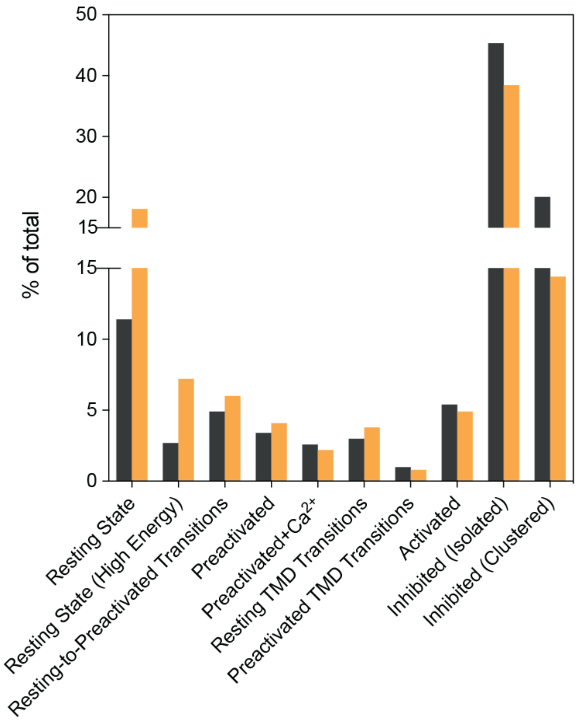

**Figure S16: Further analysis of the $Ca^{2+}$-dependent conformational landscape of $hIP_3R3$. (A)** Comparison of the relative abundances of all states from two grids prepared at 100 nM $Ca^{2+}$, shown here as black (602,609 particles) and gold (219,465 particles).

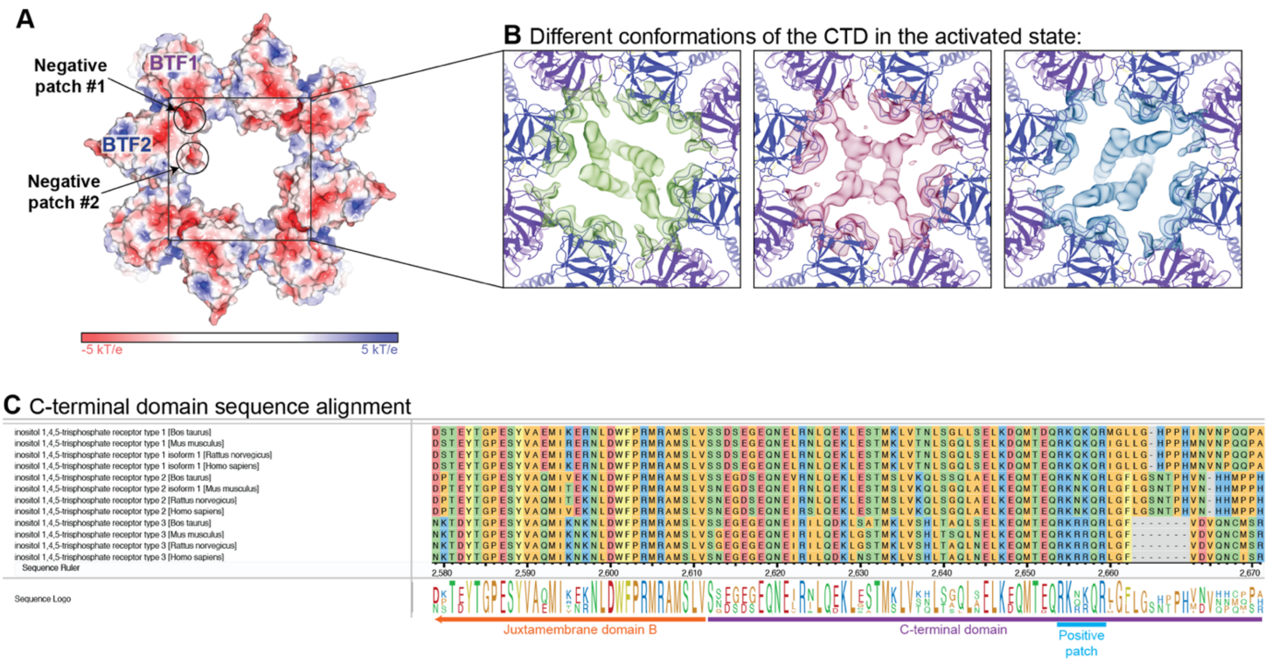

**Figure S17: The CTD dynamically interacts with two electronegative patches on the BTF ring. (A)** Electrostatic surface potential of the BTF ring of the activated state. Two electronegative patches highlighted with circles on a single protomer. **(B)** Cryo-EM density shown on the activated state model of the BTF ring for three conformations (green, red, blue). **(C)** Multiple sequence alignment shows that a stretch of positive residues on the CTD is highly conserved.

***Movies:***

**Movie M01: 6 modes of 3DVA for inhibited state.**

**Movie M02: 6 modes of 3DVA for resting state.**

**Movie M03: 6 modes of 3DVA for activated state.**

**Movie M04: 6 modes of 3DVA for preactivated state.**

**Movie M05: 6 modes of 3DVA for preactivated+$Ca^{2+}$ state.**

**Movie M06: 6 modes of 3DVA for resting-to-preactivated transitions.**

**Movie M07: 6 modes of 3DVA for resting TMD transitions.**

**Movie M08: 6 modes of 3DVA for preactivated TMD transitions.**

**Movie M09: Preactivated+$Ca^{2+}$ to activated morph, view of TMD and JD from cytosol.**

**Movie M10: Preactivated+$Ca^{2+}$ to activated morph, view of TMD and JD from membrane plane.**

**Movie M11: Preactivated+$Ca^{2+}$ to activated morph, view of TMD and JD for a single chain from membrane plane.**

***Tables:***

**Table 1: Acquisition parameters for each data set**

**Table 2: Validation of models built from density-modified composite maps**

**Table 3: Validation of models built from single density-modified maps**

**Table 4: Validation of cryo-EM density presented in Figure 3 and Figure S13**

**Table 5: Validation of cryo-EM density presented in Figure 4 and Figure S15**

**Table 6: Validation of cryo-EM density presented in Figure S17**

**Table 7: Raw particle counts for titration analysis**

**Table 8: Cal-520-AM calcium imaging analysis**

### Table 1: Acquisition parameters for each data set

| | **1 nM** | 10 nM | **100 nM** (#1) | **100 nM** (#2) | **1 µM** | **10 µM** |
|---|---|---|---|---|---|---|
| **Total Micrographs** | 637 | 2150 | 6126 | 4312 | 1327 | 3136 |
| **Microscope** | FEI Krios | FEI Krios | FEI Krios | FEI Krios | FEI Krios | FEI Krios |
| **Voltage** $(kV)$ | 300 | 300 | 300 | 300 | 300 | 300 |
| **Magnification** | 29,000 | 29,000 | 29,000 | 29,000 | 29,000 | 29,000 |
| **Dose Rate** $(e^-/pixel/sec)$ | 15 | 15 | 15 | 15 | 15 | 15 |
| **Detector** | Gatan K3 | Gatan K3 | Gatan K3 | Gatan K3 | Gatan K3 | Gatan K3 |
| **Pixel Size** $(\mathring{A})$ | 0.826 | 0.826 | 0.826 | 0.826 | 0.826 | 0.826 |
| **Exposure Time** $(sec)$ | 3 | 3 | 3 | 3 | 3 | 3 |
| **Frames** | 60 | 60 | 60 | 60 | 60 | 60 |
| **Exposure** $(e^-/\mathring{A}^2)$ | 66 | 66 | 66 | 66 | 66 | 66 |
| **Defocus Range** $(\mu m)$ | -0.5 to -2.5 | -0.5 to -2.5 | -0.5 to -2.5 | -0.5 to -2.5 | -0.5 to -2.5 | -0.5 to -2.5 |
| **Average Particles Per Micrograph Picked** | 119 | 125 | 135 | 77 | 99 | 105 |
| **Average Particles Per Micrograph Kept** | 89 | 95 | 100 | 52 | 79 | 81 |
| **Particles Kept** (%) | 75% | 76% | 74% | 68% | 80% | 77% |
| **Particles Kept (total)** | 56,681 | 203,838 | 614,301 | 225,190 | 108,079 | 254,044 |
| **Box Size** (pixels) | 512 | 512 | 512 | 512 | 512 | 512 |

### Table 2: Validation of models built from density-modified composite maps

| | Resting | Preactivated | Preactivated w/ Ca2+ | Activated | Inhibited |
|---|---|---|---|---|---|
| **EMDB ID** | EMD-XXXXX | EMD-XXXXX | EMD-XXXXX | EMD-XXXXX | EMD-XXXXX |
| ***Map Validation:*** | | | | | |
| **Resolution** (Half-map FSC = 0.143; Å) Consensus / Protomer / CD / TMD+JD / CLD+ARM3 / BTF1+BTF2+ARM1 / ARM2 | 2.7 / 2.7 / 2.7 / 2.5 / 2.8 / 2.7 / 3.3 | 3.7 / 3.7 / 3.8 / 3.6 / 4.0 / 3.9 / 6.5 | 3.6 / 3.5 / 3.4 / 3.5 / 3.5 / 3.3 / 4.2 | 3.2 / 3.1 / 3.0 / 3.0 / 3.0 / 2.9 / 3.3 | 2.5 / 2.6 / 3.0 / 2.5 / 2.7 / 3.4 / 3.4 |
| **B-factor** (Guinier plot) Consensus / Protomer / CD / TMD+JD / CLD+ARM3 / BTF1+BTF2+ARM1 / ARM2 | 81 / 84 / 88 / 81 / 91 / 89 / 120 | 68 / 93 / 121 / 115 / 125 / 147 / 802 | 67 / 80 / 89 / 117 / 109 / 100 / 146 | 80 / 85 / 86 / 98 / 95 / 86 / 125 | 80 / 90 / 122 / 105 / 105 / 179 |
| **Refinement Symmetry** | C1 | C1 | C1 | C1 | C1 |
| **Total Symmetry Expanded Particles** (Final Map) | 767,556 (271,434 for ARM2) | 186,210 | 123,416 | 228,188 | 3,668,312 (357,163 for BTF1+BTF2+ARM1) |
| **PDB ID** | XXXX | XXXX | XXXX | XXXX | XXXX |
| ***Model Composition:*** | | | | | |
| **Atoms** (Non-hydrogen) | 73,580 | 72,316 | 72,320 | 73,140 | 72,296 |
| **Chains** | 4 | 4 | 4 | 4 | 4 |
| **Protein Residues** | 9,072 | 8,996 | 8,996 | 9,088 | 8,840 |
| **Ions** Ca2+ / Zn2+ | 0 / 4 | 0 / 4 | 4 / 4 | 4 / 4 | 8 / 4 |
| **Ligands** ATP / IP3 | 4 / 4 | 4 / 4 | 4 / 4 | 4 / 4 | 4 / 4 |
| **Waters** | 608 | 0 | 0 | 0 | 644 |
| ***Model Validation:*** | | | | | |
| **Map-to-Model** (FSC = 0.5 for density-modified composite of local refinements) | 2.77 | 3.87 | 3.61 | 3.05 | 2.62 |
| **Map-to-Model** (CC) BTF1 / BTF2 / ARM1 / CLD / ARM2 / ARM3 / JD / TMD / CTD | 0.82 / 0.79 / 0.78 / 0.76 / 0.76 / 0.77 / 0.79 / 0.76 / 0.54 | 0.75 / 0.73 / 0.72 / 0.75 / 0.74 / 0.75 / 0.75 / 0.73 / 0.66 | 0.82 / 0.82 / 0.81 / 0.81 / 0.80 / 0.78 / 0.79 / 0.76 / 0.64 | 0.74 / 0.73 / 0.74 / 0.74 / 0.75 / 0.74 / 0.72 / 0.71 / 0.56 | 0.28 / 0.49 / 0.72 / 0.80 / 0.73 / 0.83 / 0.83 / 0.80 / 0.70 |
| **Mean B-Factor** $(\AA^2)$ Protein / Ligand / Water (NA if not present) | 87.8 / 99.2 / 67.0 | 89.3 / 79.5 / NA | 78.3 / 75.7 / NA | 119.1 / 118.3 | 93.9 / 100.0 / 57.4 |
| **R.M.S. Deviations** Bond Lengths (Å) / Bond Angles (°) | 0.002 / 0.447 | 0.002 / 0. 449 | 0.002 / 0.428 | 0.002 / 0.463 | 0.002 / 0.409 |
| **MolProbity Score** | 1.14 | 1.37 | 1.32 | 1.21 | 1.20 |
| **CaBLAM outliers** (%) | 1.00 | 1.10 | 1.19 | 1.27 | 0.70 |
| **Clash Score** | 3.45 | 6.78 | 5.84 | 4.24 | 4.18 |
| **Rotamer Outliers** (%) | 0.30 | 0.10 | 0.10 | 0.49 | 0.20 |
| **Cβ Deviations** | 0.00 | 0.00 | 0.00 | 0.00 | 0.00 |
| **Ramachandran Plot** (%) Favored / Allowed / Disallowed | 98.39 / 1.61 / 0.00 | 98.02 / 1.98 / 0.00 | 0.002 / 0.428 | 98.17 / 1.83 / 0.00 | 98.21 / 1.79 / 0.00 |

### Table 3: Validation of models built from single density-modified maps

| | Higher-Order Inhibited Assembly | Labile Resting #1 | Labile Resting #2 |
|---|---|---|---|
| EMDB ID | EMD-XXXXX | EMD-XXXXX | EMD-XXXXX |
| ***Map Validation:*** | | | |
| **Resolution (Half-map FSC = 0.143; Å)** | 3.4 | 3.5 | 3.6 |
| **B-factor (Guinier plot)** | 88 | 53 | 57 |
| Refinement Symmetry | C1 | C1 | C1 |
| Total Symmetry Expanded Particles (Final Map) | 78,512 | 91,326 | 155,671 |
| PDB ID | | | |
| ***Model Composition:*** | | | |
| Atoms (Non-hydrogen) | 11,906 | 79,972 | 79,972 |
| Chains | 2 | 4 | 4 |
| Protein Residues | 1,471 | 9,072 | 9,072 |
| Ions Ca2+ / Zn2+ | 2 / 0 | 0 / 4 | 0 / 4 |
| Ligands ATP / IP3 | 0 / 0 | 4 / 4 | 4 /4 |
| Waters | 0 | 0 | 0 |
| ***Model Validation:*** | | | |
| **Map-to-Model (FSC = 0.5 for density-modified composite of local refinements)** | 3.83 | 3.61 | 3.88 |
| Map-to-Model (CC) BTF1 / BTF2 / ARM1 / CLD / ARM2 / ARM3 / JD / TMD / CTD | - / - / - /0.70/0.73/ - / - / - / - | 0.80/0.77/0.78/0.71/0.63/0.75/0.78/0.75/0.44 | 0.76/0.75/0.75/0.70/0.61/0.70/0.72/0.67/0.41 |
| Mean B-Factor ($\mathring{A}^2$) Protein / Ligand / Water (NA if not present) | 118.1 / 148.5 | 79.5 / 59.8 | 54.1 / 45.7 |
| R.M.S. deviations Bond Lengths (Å) / Bond Angles (°) | 0.002 / 0.437 | 0.002 / 0.443 | 0.002 / 0.472 |
| MolProbity Score | 1.30 | 1.26 | 1.35 |
| CaBLAM outliers (%) | 0.42 | 0.72 | 1.00 |
| Clash Score | 5.56 | 4.91 | 6.31 |
| Rotamer Outliers (%) | 0.00 | 0.10 | 0.15 |
| Cβ Deviations | 0.00 | 0.00 | 0.00 |
| Ramachandran Plot (%) Favored / Allowed / Disallowed | 98.76 / 1.24 / 0.00 | 98.75 / 1.25 / 0.00 | 98.39 / 1.61 / 0.00 |

### Table 4: Validation of cryo-EM density presented in Figure 3 and Figure S13

| | Figure 3A (3/1 ext/ret) | Figure 3A (2/2 ext/ret) | Figure 3A (1/3 ext/ret) | Figure 3B | Figure 3C | Figure 3D | Figure 3E | Figure 3F | Figure 3G |
|---|---|---|---|---|---|---|---|---|---|
| EMDB ID | EMD-XXXXX | EMD-XXXXX | EMD-XXXXX | EMD-XXXXX | EMD-XXXXX | EMD-XXXXX | EMD-XXXXX | EMD-XXXXX | EMD-XXXXX |
| **Resolution (Half-map FSC = 0.143; Å)** | 4.0 | 4.0 | 3.9 | 4.0 | 3.7 | 4.1 | 3.8 | 3.7 | 4.0 |
| **B-factor (Guinier Plot)** | 52 | 24 | 17 | 48 | 69 | 90 | 97 | 82 | 52 |
| Low-Pass Filter (Å) | 5 | 5 | 5 | 5 | 5 | 5 | 5 | 5 | 5 |
| Refinement Symmetry | C1 | C1 | C1 | C1 | C1 | C1 | C1 | C1 | C1 |
| Total Symmetry Expanded Particles (Final Map) | 83,868 | 31,541 | 25,729 | 21,877 | 62,262 | 87,974 | 100,749 | 81,743 | 24,087 |

***Note:** Figure 3A maps are representative and not exhaustive i.e. multiple classes produced these conformations that were later combined into the resting-to-preactivated transition ensemble.

### Table 5: Validation of cryo-EM density presented in Figure 4 and Figure S15

| | Preactivated TMD Transition (−C2) | Preactivated TMD Transition (−C4) | Resting TMD Transition (−C2) | Resting TMD Transition (−C4) |
|---|---|---|---|---|
| EMDB ID | EMD-XXXXX | EMD-XXXXX | EMD-XXXXX | EMD-XXXXX |
| **Resolution (Half-map FSC = 0.143; Å)** | 3.7 | 3.6 | 3.6 | 3.7 |
| **B-factor (Guinier Plot)** | 33 | 44 | 35 | 46 |
| Low-Pass Filter (Å) Overview / Zoomed | 4 / 7 | 4 / 7 | 4 / 7 | 4 / 7 |
| Refinement Symmetry | C1 | C1 | C1 | C1 |
| Total Symmetry Expanded Particles (Final Map) | 34,006 | 66,872 | 63,075 | 121,444 |

***Note:** Maps from Figure 4 and Figure S15 not presented here can be found on Table 2.

### Table 6: Validation of cryo-EM density presented in Figure S17

| | Figure S13J | Figure S13K | Figure S13L | Figure S13M | Figure S13N | Figure S13O |
|---|---|---|---|---|---|---|
| EMDB ID | EMD-XXXXX | EMD-XXXXX | EMD-XXXXX | EMD-XXXXX | EMD-XXXXX | EMD-XXXXX |
| **Resolution (Half-map FSC = 0.143; Å)** | 3.8 | 3.1 | 3.1 | 3.5 | 3.2 | 3.0 |
| **B-factor (Guinier Plot)** | 103 | 58 | 81 | 105 | 88 | 67 |
| Refinement Symmetry | C1 | C1 | C1 | C1 | C1 | C1 |
| Total Symmetry Expanded Particles (Final Map) | 195,763 | 49,313 | 176,457 | 194,283 | 186,458 | 89,264 |

### Table 7: Raw particle counts for titration analysis

| | 1 nM | 10 nM | 100 nM (#1) | 100 nM (#2) | 1 µM | 10 µM |
|---|---|---|---|---|---|---|
| Resting State | 25248 | 54108 | 68808 | 39701 | 2296 | 1728 |
| Labile Resting States | 11416 | 13412 | 16435 | 15877 | 1605 | 3005 |
| Resting TMD Transitions | 3353 | 7663 | 18093 | 8247 | 2706 | 6070 |
| Resting-to-Preactivated Transitions | 4182 | 21176 | 29757 | 13212 | 1096 | 1252 |
| Preactivated | 2013 | 12804 | 20482 | 8929 | 870 | 1454 |
| Preactivated (+Ca2+) | 615 | 8896 | 15643 | 4792 | 520 | 388 |
| Preactivated TMD Transitions | 184 | 2051 | 6131 | 1862 | 437 | 222 |
| Activated | 565 | 8218 | 32272 | 10684 | 2978 | 2330 |
| Inhibited (Clustered) | 3322 | 24419 | 121239 | 31703 | 18529 | 47031 |
| Inhibited (Isolated) | 4988 | 48075 | 273749 | 84458 | 74422 | 185143 |

**Note:** Non-symmetry expanded particle counts.

### Table 8: Cal-520-AM calcium imaging analysis

| Cell line/construct | IP3R-null | hIP3R3 | CD site mutant | JD site mutant | JD/CD sites double mutant | dARM2 | d(S1'-S1") | BTF ring single mutant | BTF ring double mutant | dCTD |
|---|---|---|---|---|---|---|---|---|---|---|
| Tag (w/ Position at N or C terminus) | - | N-10xHis-EGFP | N-10xHis-EGFP | N-10xHis-EGFP | N-10xHis-EGFP | N-10xHis-EGFP | N-10xHis-EGFP | N-10xHis-EGFP | N-10xHis-EGFP | N-10xHis-EGFP |
| Linker Sequence (Tag-Linker-Gene) | - | LEVLFQGPSRV | LEVLFQGPSRV | LEVLFQGPSRV | LEVLFQGPSRV | LEVLFQGPSRV | LEVLFQGPSRV | LEVLFQGPSRV | LEVLFQGPSRV | LEVLFQGPSRV |
| Mutation | - | - | E1125Q | E1882Q, E1946Q | E1882Q, E1946Q, E1125Q | Δ(A1101-W1586) | Δ(D2252-I2302) | K169A | W168A, K169A | Stop codons at L2629x, N2630x |
| Mean inter-spike interval (ISI) (seconds) | - | 21.7 | 12.7 | - | - | 117.1 | 17.6 | - | - | 12.1 |
| Lower 95% CI of mean (ISI) (seconds) | - | 20.9 | 12.4 | - | - | 81.0 | 16.5 | - | - | 11.7 |
| Upper 95% CI of mean (ISI) (seconds) | - | 22.5 | 13.0 | - | - | 153.2 | 18.7 | - | - | 12.4 |
| Median no. of peaks/cell | - | 4 | 6 | - | - | 1 | 3 | - | - | 4 |
| Normalized fluorescence/second at half maxima ± SEM (n=3) | - | 0.103 ± 0.015 | 0.143 ± 0.011 | 0.028 ± 0.006 | 0.033 ± 0.003 | 0.173 ± 0.010 | 0.130 ± 0.019 | 0.026 ± 0.001 | - | 0.188 ± 0.013 |
